# Complete Mutagenesis of the Genome and Proteome of ΦX174

**DOI:** 10.64898/2026.07.25.740675

**Authors:** Huijin Wei, Xianghua Li, Ben Lehner

## Abstract

The bacteriophage ΦX174 was the first genome to be sequenced and the first to be chemically synthesised. Here we present a complete map of the consequences of changing every nucleotide in the ΦX174 genome and every amino acid in the proteome. In total, half of nucleotide and >60% of amino acid changes impair fitness. However, this varies substantially across proteins and non-coding regions. Surprisingly, the most important mechanistic cause of reduced fitness is disruption of protein interaction interfaces, accounting for nearly half of detrimental protein variants. A further one quarter of damaging variants disrupt protein cores but one in four lack a mechanistic explanation. Mutational effects are only moderately-well predicted by state-of-the-art artificial intelligence models, but, combined with structural modelling, provide mechanistic hypotheses and insights into the DNA replication machinery. This complete mutagenesis of a biological system quantifies the limits of our current understanding of, and ability to predict, molecular biology.

## Introduction

The genomics era was initiated in 1977 with the sequencing of the genome of the bacteria-infecting virus ΦX174 (family *Microviridae*) (*1*). A quarter of a century later, ΦX174 also became the first genome to be completely chemically synthesised, heralding the era of synthetic genomics (*2*). Isolated in the 1930s (*3*), ΦX174 was the first virus identified with a ssDNA genome (*4*) and the first shown to encode proteins in overlapping open reading frames (ORFs) (*5*).

We reasoned that ‘complete mutagenesis’ of ΦX174 - the comprehensive mutagenesis of all of its molecular components - would be a good test of both our current mechanistic understanding of biology and of our ability to predict how a biological system responds to perturbations. ΦX174 is a very extensively studied and well-defined system. The 5,386 nucleotide (nt) circular genome encodes six structural proteins (B, D, F, G, H, and J) involved in capsid assembly, as well as five non-structural proteins (A, Astar, C, E, and K) that mediate viral genome replication, synthesis, packaging and host cell lysis (*6*). Many experimental structures have been determined (*7–10*) and the virus has served as a key model system for more than 50 years (*11–13*). The ΦX174 life cycle starts with host recognition and attachment, followed by genome ejection into the host cell. The ejected ssDNA is converted into replicative form double-stranded DNA (dsDNA), enabling rolling-circle replication, transcription, viral protein expression, and virion capsid assembly (*6*) (Fig. 1a).

**Figure 1:**
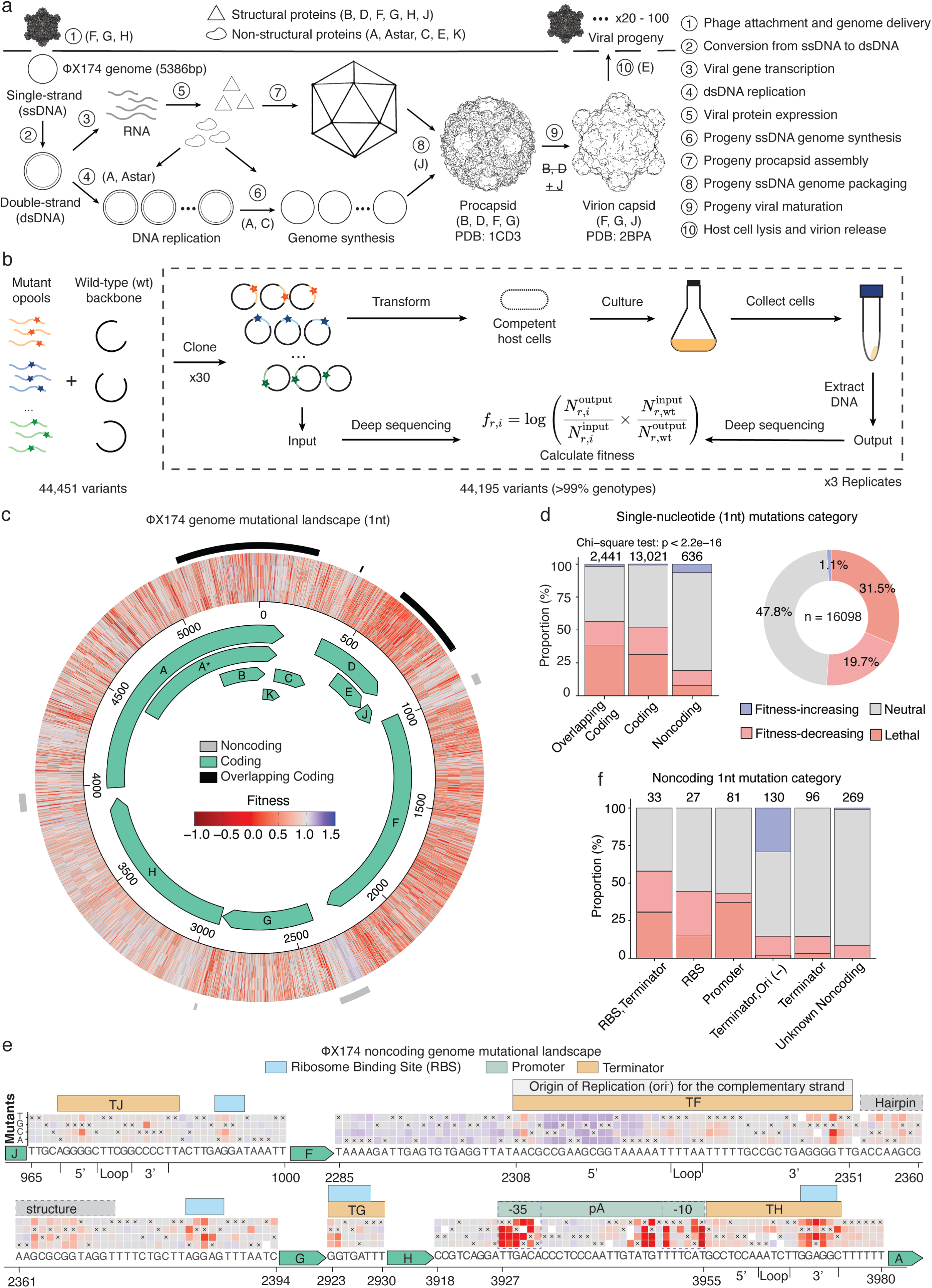
Complete mutagenesis of ΦX174 genome and proteome. a, Overview of the viral life cycle. b, Schematic of the experimental design. c, Genome-wide fitness landscape at single-nucleotide resolution. Circular bars indicate genomic regions, including noncoding regions (gray) and overlapping coding regions (black). Open reading frames (ORFs) are shown as circular arrow bars inside the heatmap. d, Mutation categories for single-nucleotide substitution (1nt) variants across genomic regions. e, Heatmaps showing the fitness effects of single-nucleotide (1nt) substitutions in noncoding regions. Bars above the heatmaps indicate annotated regulatory elements, including ribosome-binding sites (RBSs), promoters, terminators, and the origin of replication for the complementary (-) strand. The 5′ stem, loop, and 3′ stem of hairpin structures are labelled below the axis. f, Mutation categories for single-nucleotide substitution (1nt) variants in noncoding regions, across annotated regulatory elements.

Our goal in mutating the entire genome and proteome (*14*, *15*) was fourfold: [1] to provide a complete description of the effects on fitness of all possible nt and amino acid (aa) substitutions in a genome and proteome; [2] to quantify the importance of different molecular mechanisms by which mutations impair fitness in a system where all measurements are made on the same biologically meaningful scale, viral fitness; [3] to test our current mechanistic understanding of a very well-studied biological system; and [4] to evaluate the predictive performance of state-of-the-art artificial intelligence (AI) models using a large and well-calibrated out-of-distribution experimental dataset. We believe that this complete mutagenesis of a biological system is quite a general approach to evaluate the limits of our current understanding of, and ability to predict, molecular biology.

## Results

### Complete mutagenesis of a genome and proteome

To quantify the effects of every nt substitution in the genome of ΦX174 and every aa substitution in its proteome, we constructed 30 overlapping oligonucleotide libraries each covering a 200-300nt fragment (Fig. S1a-c). Each variant library was assembled into the viral genome, and the fitness of variants quantified in the native *Escherichia coli* host after 80min (2-3 generations) (Fig. 1b).

In total, we quantified the fitness effects of 44,195 variants, representing >99% of designed genotypes. This includes 16,098 single-nucleotide (1nt) variants, 40,358 amino-acid variants, and 1,396 synonymous variants. Fitness measurements were highly reproducible across three biological replicates (pairwise Pearson’s r=0.988-0.992; median r=0.99; Fig. S1d). 8,229 of 16,098 (51.2%) single nucleotide variants reduced fitness (false discovery rate, FDR < 0.05, Benjamini–Hochberg correction) (Fig. 1c). Of these, 5,065 (62%) caused a greater than 50% reduction in fitness and are hereafter referred to as lethal variants. Overall, 68.5% (3,690/5,386) of nucleotide positions in the genome are sensitive to mutation, with nearly half (48%, 2,588/5,386) of nucleotide positions classified as essential based on the presence of at least one lethal mutation (Fig. S1i).

The proportion of detrimental single nucleotide substitutions varies across genomic regions (Fig. 1c-d). Mutations in overlapping protein-coding regions, which comprises 15.1% of the genome, are more likely to reduce fitness, with 56.3% of mutations (1,374/2,441) having a negative impact compared to 51.7% in single protein-coding regions (6,733/13,021, p=3.28×10⁻^5^, Fisher’s exact test (FET)). In contrast, only 19.2% of mutations (122/636) in the noncoding regions reduce fitness. Overlapping protein-coding regions also contain the highest proportion of essential nucleotides (59%), followed by other coding regions (47.9%) and noncoding regions (11.1%) (Fig. S1i).

### The non-coding genome

The four non-coding regions of ΦX174 comprise 4% of the genome (217nt) and are densely packed with regulatory elements, including four ribosome binding sites (RBS), one promoter (*pA)*, four transcription terminators (T_J_, T_F_, T_G_, T_H_), and the origin of replication (ori^−^) for the complementary (-) strand (*1*, *16*, *17*) (Fig. 1e). A majority of detrimental noncoding mutations (81.1%, 99/122) occur within annotated regulatory elements (Fig. 1e-f). Specifically, 31.1% (38/122) are found within regulatory elements with overlapping annotations, such as terminators with RBSs or ori^−^. 28.7% (35/122) are found in the pA promoter, 11.5% (14/122) in terminators, and 10.0% (12/122) in RBSs. Notably, a higher proportion of mutations in the promoter (43.2%, 35/81) are detrimental compared to those in terminators (14.6%, 14/96) (Fig. 1f).

Analysis of the pA promoter reveals that 15 of 35 detrimental mutations localised to the −35 element (TTGACA, nt 3927-3932), while 11 mapped to a degenerate -10 element (TTTCAT, nt 3950-3955) (*18*) with an upstream motif extension (TATG, nt 3945-3948) that corresponds to the TRTG motif, a binding site for the σ⁷⁰ RNA polymerase holoenzyme (*19*). Notably, 9 out of 10 mutations in the TATG motif are detrimental, compared to 11 out of 18 in the -10 element, highlighting the essential function of this motif extension (Fig. 1e). Notably, the only tolerated mutation, A3946G, also belongs to the TRTG motif, which has been reported to enhance holoenzyme binding in bacteria (*20*).

All four terminators (T_J,_ T_F,_ T_G,_ and T_H_) in the non-coding genome are Rho-independent and rely on stem-and-loop hairpin secondary structures for transcription termination (*16*). However, mutational sensitivity across pairing stems in two terminators, T_F_ and T_H_, is asymmetric (Fig. 1e), likely due to disruption of overlapping regulatory elements embedded in only one stem. For example, in the T_H_ terminator hairpin (nt 3967-3975), 63% (17/27) of detrimental mutations are located in the 3′ stem, which overlaps with RBS of gene A (GGAGG, nt 3969-3973), compared to 13% (3/23) in the 5’ stem.

Mutational effects can also have opposing directions when regulatory elements overlap. In the T_F_ terminator that overlaps with ori^−^, 39 mutations in the 5’ stem (nt 2308-2326) increase fitness, whereas mutations in the 3’ stem and loop tend to be neutral or to decrease fitness (Fig. 1e). This hairpin is also a binding site for PriA, an essential 3’-5’ DNA helicase for the phage DNA replication, which binds DNA substrates in a structure-specific manner (*17*, *21*). The asymmetric mutational pattern reflects the polarity of PriA binding or function. Many fitness-increasing mutations in the 5′ stem are predicted to disrupt local base pairing that may correspond to a more accessible structural feature for PriA binding (*21*). Consistent with this hypothesis, a mutation in the 5’ stem (T2321C) that is predicted to establish base-pairing with a 3’ stem residue (G2338) does not improve fitness (Fig. S1j).

In addition, a shorter hairpin located immediately downstream of the T_F_ terminator (nt 2353-2373) shows a more symmetric mutational pattern. This region lies upstream of a poly-U tract and is predicted to overlap part of a putative Rho-independent terminator (*16*), containing 16 detrimental mutations that are distributed across both stems. This suggests the shorter hairpin structure may be functionally important and likely contributes to transcription termination together with the annotated T_F_ terminator.

### Mutational landscape of the proteome

We obtained fitness measurements for 99.5% (39,719/39,920) of all possible single amino acid mutations across the proteome, comprising 29,979 single-ORF mutations and 9,740 multi-ORF mutations that alter the sequence of more than one protein. Multi-ORF mutations are excluded from all subsequent analyses unless otherwise stated.

Nearly all nonsense mutations (99%, 1,492/1,507) are lethal (Fig. 2a). Nonsense variants in protein K are the exception, indicating this protein is not required for fitness in the tested conditions. 76% of stop-loss mutations (72/92) are detrimental. Protein F is an exception, with 16 stop-loss mutations tolerated, likely due to the presence of an in-frame stop codon located immediately downstream of the native stop codon.

**Figure 2:**
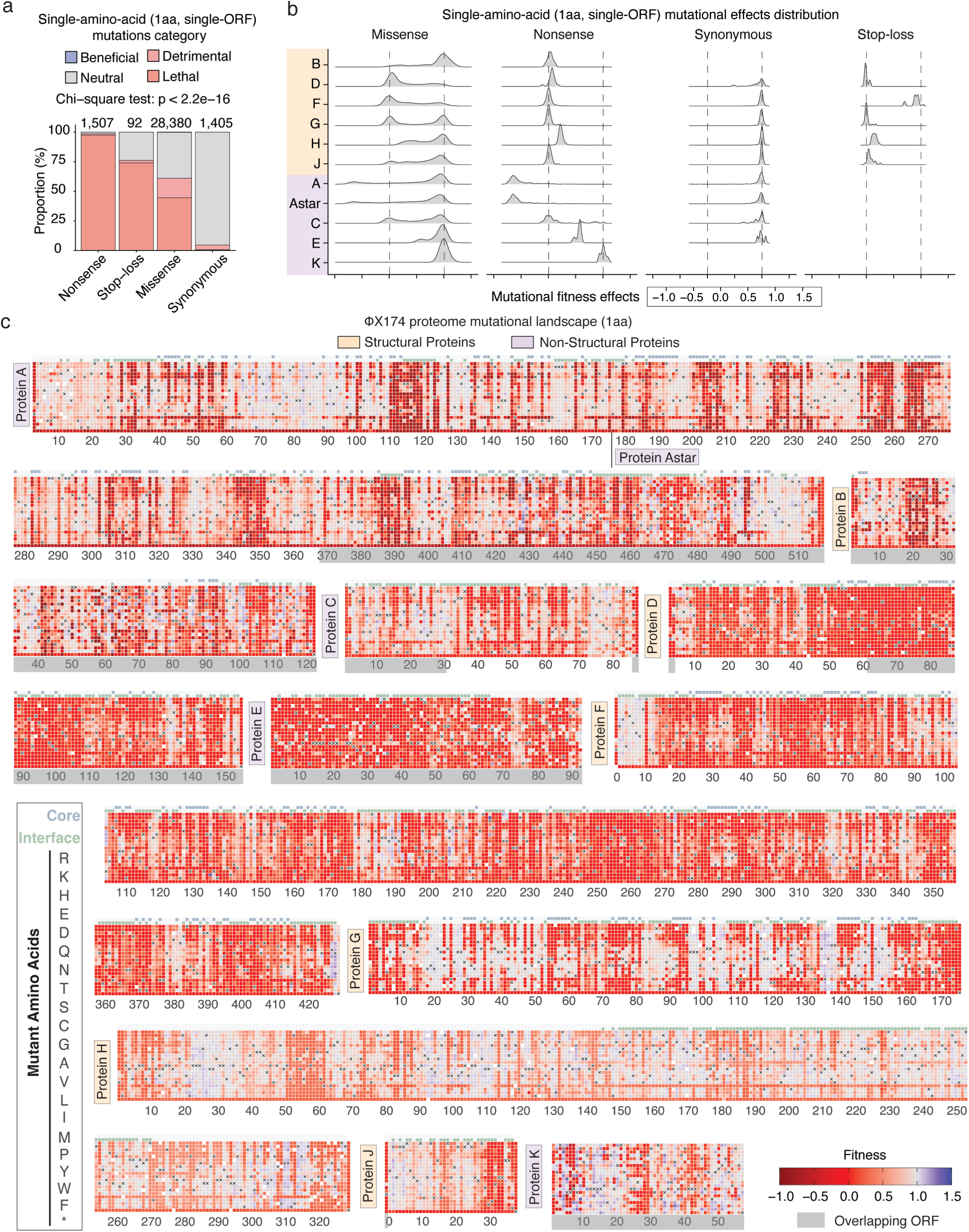
Complete mutagenesis of the ΦX174 proteome. a, Mutation categories for single-amino-acid variants, grouped by mutation type. b, The distributions of fitness effects for single amino acid mutations (single-ORF). c, Heatmaps showing variant fitness landscapes for 11 viral proteins, proteins A and A* are shown in a shared heatmap. Additional rows above each heatmap indicate core and interface residues. Positions shaded in gray correspond to regions encoded by overlapping ORFs.

The distribution of fitness effects for single aa missense mutations is bimodal with peaks centred at the modes for nonsense and synonymous variants for each protein (Fig. 2b). In total, 70% (19,908/28,380, FDR<0.05) of missense mutations are detrimental and 44.6% (12,658/28,380, FDR<0.05) are lethal. Notably, the proportion of detrimental mutations varies substantially across different proteins (Fig. 3a). Substitutions that increase fitness are extremely rare (1.4%; 403/28,380, FDR<0.05), with the majority occurring in structural proteins (Fig. S2b), such as protein F (37%; 149/403), protein H (33%; 134/403) and protein J (11%; 45/403). This is consistent with previous studies showing that viral structural proteins are important substrates for adaptation (*14*, *15*). In addition, beneficial mutations are significantly enriched in interaction interfaces compared to neutral mutations (55.3%, 223/403 vs. 36.8%, 2973/8069; Odds ratio (OR) =2.1, FET p=2.19×10⁻^13^) (Fig. S2c).

**Figure 3:**
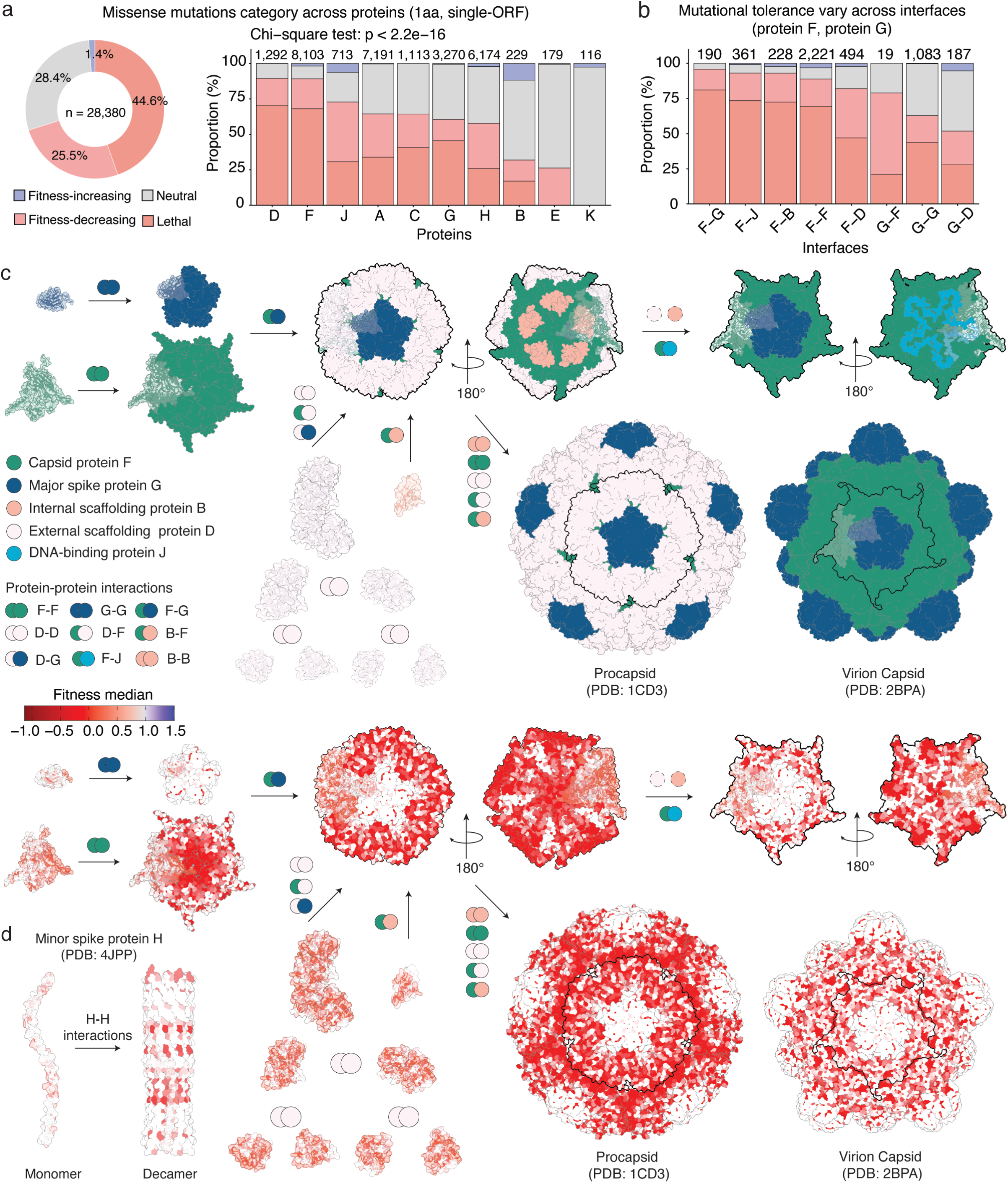
Mutational tolerance varies across proteins and interfaces. a, Mutation categories for single amino-acid substitutions (single-ORF, missense mutations) across the proteome and individual viral proteins. b, Mutation categories across interfaces in the capsid protein F and major spike protein G. c, Protein structures showing procapsid assembly and virion maturation. The viral procapsid assembles in a defined order from capsid protein F, major spike protein G, external scaffolding protein D, and internal scaffolding protein B. During maturation, scaffolding proteins B and D dissociate, and DNA-binding protein J binds capsid protein F as the viral genome is packaged. Protein structures in the lower schematics are colored by positional median fitness effect. The procapsid structure is from PDB: 1CD3, and the virion capsid structure is from PDB: 2BPA. Assembly-intermediate pentamer structures are shown after a 180° clockwise rotation to display internal surfaces. d, Assembly of the decameric DNA-transport tunnel formed by minor spike protein H, showing the resolved coiled-coil region from PDB: 4JPP colored by positional mediant fitness effects.

### Synonymous variants

Most synonymous mutations are neutral (92%; 1289/1,405) (Fig. 2a). Among 116 detrimental synonymous mutations, only 23 overlap known regulatory elements. These include nine detrimental synonymous mutations in protein A (Q107, L108, D109, I110, N111, N112, T113, I114, and D115) that colocalise with the origin of replication for the viral (+) strand (ori^+^) (*22*). Four synonymous mutations in protein A (L295, S301, Y303, S304) and six in protein C (V63, D64, L71, L72, S73, S75) colocalise with the promoters *pB* and *pD*, respectively (*1*, *16*). Two synonymous mutations in protein C (I82, G83) and one in protein D (E56) colocalise with RBS for the expression of their downstream proteins D and E (*1*). Additionally, one synonymous mutation in protein G (P173) colocalise with the terminator T_G_ (*16*). However, most detrimental synonymous mutations (93/116) remain mechanistically unexplained, although they may affect fitness through altered codon usage (*23*). When clustered and integrated with features predicted from sequences, these mutations can also identify potential regulatory elements (*16*). For example, thirteen synonymous mutations in protein A (R271, T272, L273, P274, T275, S277, V278, D279, P280, F282, G283, R284, R287) map to a previously predicted promoter region (*1*), suggesting that this promoter is also functionally important.

### Mutational tolerance varies across proteins

External scaffolding protein D (89.4% detrimental, 1,155/1,292) and capsid protein F (89.1% detrimental, 7,222/8,103) are the least tolerant of mutations (Fig. 2c, 3a). Protein D mediates morphogenesis, forms the primary contacts during procapsid assembly, and appears to be highly evolutionarily constrained (*7*, *24*, *25*), whereas protein F functions as an interaction hub, interacting with other structural proteins during assembly, maturation, and infection (*7*, *8*, *24*). Among the remaining viral procapsid and virion capsid proteins, detrimental mutations accounted for 31.9% to 72.8% of variants, a range comparable to that reported in previous phage protein mutagenesis studies (*26*, *27*). Specifically, DNA-binding protein J exhibits the highest proportion (72.8%; 519/713), followed by major spike protein G (60.5%; 1,977/3,270), minor spike protein H (57.8%; 3,569/6,174), and internal scaffolding protein B (31.9%; 73/229). Protein B supports procapsid assembly from the interior and is replaced by DNA-binding protein J during the viral maturation (*8*). Spike proteins G and H mediate host attachment, followed by genome delivery through a tunnel formed by decameric protein H (*9*, *28*).

Among non-structural proteins, only replication-associated proteins A (64.4%; 4,630/7,191) and C (64.3%; 716/1,113) show comparable proportions of detrimental mutations as structural proteins (Fig. 3a). Protein A is a multifunctional enzyme that cleaves and ligates the viral (+) strand during viral DNA replication (*29*). Protein C is an inhibitor of DNA replication initiation and is required for ssDNA genome synthesis and packaging (*30*). Notably, detrimental mutations in protein A are associated with larger fitness reductions than mutations in other proteins, likely reflecting direct roles in DNA replication and viral genome amplification within host cells.

The remaining non-structural proteins contain far fewer detrimental mutations. Only 26.3% (47/179) of mutations in lysis protein E, which inhibits the host cell peptidoglycan biosynthesis and promotes host cell lysis (*10*), are detrimental. Protein K has been reported to modulate burst size (*31*), but no mutations in protein K impair viral fitness in our selection assay (Fig. 3a). However, analyses of these two proteins are limited because nearly 90% of mutations also affect other proteins (1,532/1,711 in protein E and 948/1,064 in protein K) and are excluded from the analysis.

In total, 79,7% (1,584/1,986) of single aa residues in the proteome are sensitive to mutations, among which 87.6% (69.8% of total, 1,387/1,986) are essential for viral fitness. The proportion of essential residues again varies across proteins (Fig. S3a), although there are important differences between residue-level and mutation-level patterns. For instance, ∼90% of mutations in protein F (7,222/8,103) are detrimental and ∼94% of residues are essential (402/427), whereas protein H shows a similarly high fraction of essential residues (81.4%; 267/328) but a lower fraction of detrimental mutations (57.8%; 3,569/6,174).

### One in four detrimental mutations disrupt protein cores

Mutations in the buried hydrophobic cores of proteins are particularly detrimental for stability (*32*, *33*). Consistent with this, 25.4% (5,058/19,908) of all detrimental mutations in the ΦX174 proteome occur at core residues (relative solvent accessibility, RSA < 25%) (Fig. 4b). Mutations in core residues are more likely to be detrimental than mutations in surface residues, with 87.5% (5,058/5,783) of core mutations classified as detrimental compared with 68.0% (12,280/18,058) of surface mutations (OR=3.3, FET p<2.2×10^−16^) (Fig. 4c). However, the enrichment of detrimental mutations in protein cores varies across proteins: 98.5% (262/266) in protein D (OR=8.3, FET p=2.9×10⁻^8^), 95.5% (2,048/2,144) in protein F (OR=3.2, FET p<2.2×10^−16^), 85.1% (2,007/2,359) in protein A (OR=4.8, FET p<2.2×10^−16^), 73.5% (731/994) in protein G (OR=2.3, FET p<2.2×10^−16^), and 50% (10/20) in Protein B (OR=1.3; FET p=0.6) (Fig. 4d).

**Figure 4:**
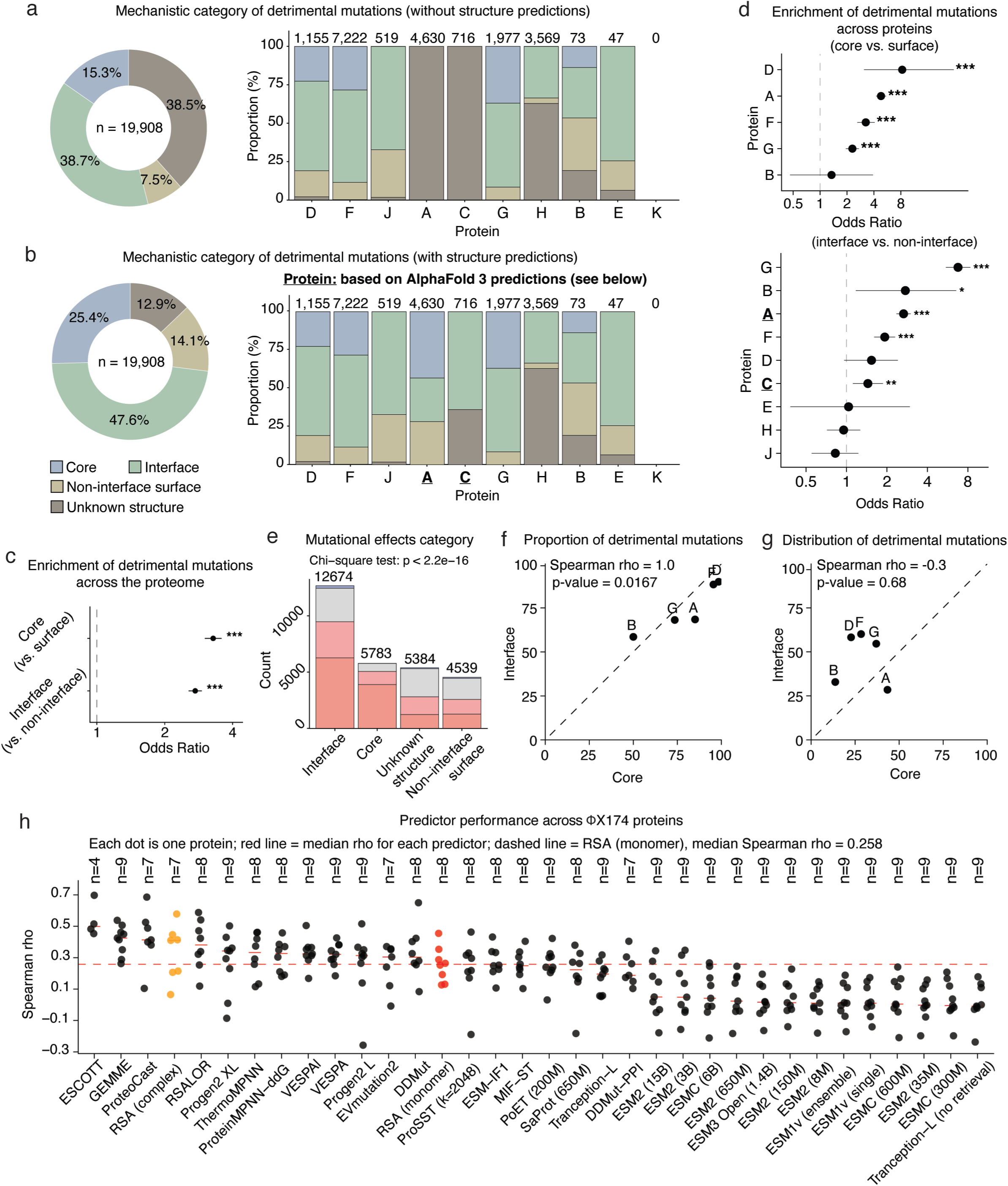
Disruption of protein cores and interaction interfaces explains 75% detrimental mutations. a,b, Mechanistic interpretation of detrimental single amino-acid substitutions across the proteome and individual viral proteins, without AlphaFold 3 structure predictions (a) and with AlphaFold 3 structure predictions (b). c-d, Enrichment of detrimental mutations in protein cores vs. surfaces and at interfaces vs. non-interface surface residues across the proteome (c) and individual viral proteins (d). Dots show odds ratios, and asterisks indicate significance from two-sided Fisher’s exact tests (*p < 0.05, **p < 0.01, ***p < 0.001). e, Counts and mutation categories across residue classes. f, The relationship between the proportion of detrimental mutations in core residues vs. interface residues across proteins. g, The relationship between the distribution of detrimental mutations assigned to core residues and interface residues across proteins. h, Evaluation of variant effect predictors. Predictors are ordered from left to right by their median performance across available proteins. Red horizontal marks indicate the median performance of each predictor, and the red dashed line indicates the median performance of the structural baseline, monomer-derived RSA. Monomer-derived RSA is shown in red, and complex-derived RSA is shown in orange.

### Nearly half of detrimental mutations are located in interaction interfaces

Beyond protein core disruption, a second hypothesised important cause of detrimental mutations is disruption of molecular interaction interfaces (defined as a minimal heavy-atom distance of less than 5 Å between proteins) (*34*, *35*). In total, 43.2% (857/1,986) of residues in the ΦX174 proteome are located in at least one interaction interface, including extensive interactions among proteins B, D, F, G, and J during viral assembly and maturation that have been well-characterized (*8*, *24*) (Fig. 3c), interactions between protein H monomers that assemble into a decameric tunnel for DNA ejection (Fig. 3d) (*9*), interactions between protein E with host membrane proteins to promote lysis (Fig. S3b), and predicted interactions between protein A with DNA and proteins A-C (Fig. 5) (*29*, *30*).

**Figure 5:**
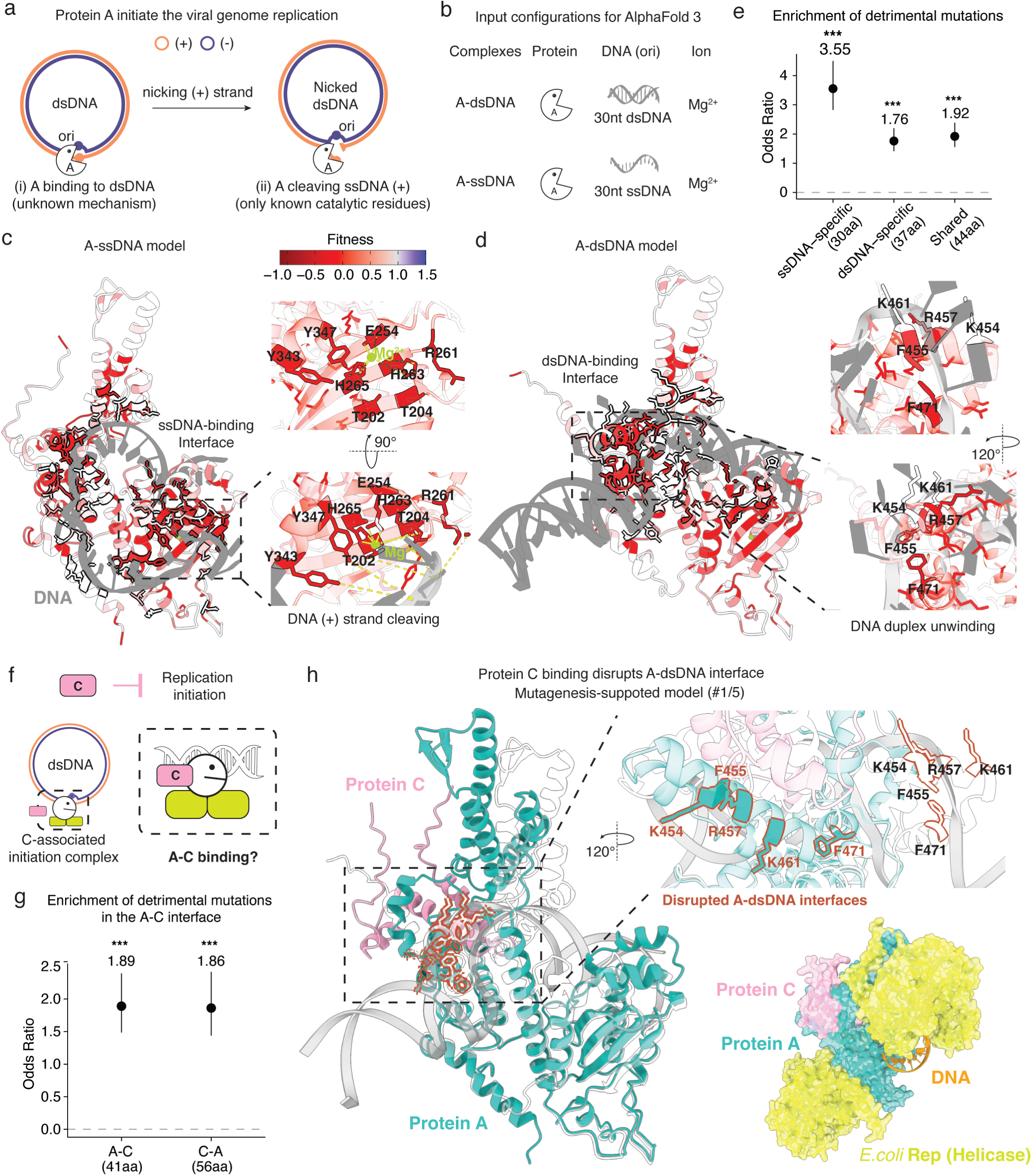
Protein C allosterically inhibits protein A-DNA interaction. a, Schematic showing two proposed steps in protein A-mediated initiation of DNA replication: binding to dsDNA followed by the cleavage of the viral (+) strand. b, Input configurations used for AlphaFold 3 structure prediction of A-dsDNA and A-ssDNA complexes; c-d, Top-ranked AlphaFold 3-predicted protein–DNA complex structures for A-ssDNA (c) and A-dsDNA (d). Protein A is colored by positional median fitness effects, and DNA is shown in gray. Protein A-DNA interfaces are outlined, and key functional residues in each interface are shown in the zoomed-in views. e, Enrichment of detrimental mutations in ssDNA-specific, dsDNA-specific, and shared A-DNA interfaces. f, Schematic showing the current understanding of protein C-mediated inhibition of DNA replication initiation. g, Enrichment of detrimental mutations at the predicted A-C interfaces in the mutagenesis-supported model with the highest A-C interface ipTM score. h, Structural model showing that protein C binds protein A near the A-dsDNA interface and disrupts the dsDNA-specific interface predicted to promote local DNA unwinding. The complete complex structure is shown at the bottom right.

Strikingly, nearly half the detrimental mutations are located in an interface (47.6%, 9,478/19,908). Interface mutations are more likely to be detrimental than mutations at non-interface surface residues (9,478/12,674, 74.8% vs. 2,802/5,384, 52%; OR=2.73, FET p<2.2×10^−16^) (Fig. 4c). Mutational tolerance varies substantially across interfaces, both when comparing the interfaces of different proteins and when comparing different interfaces in the same protein (Fig. 3b, Fig. S3c). Comparing the interfaces of different proteins, the interfaces of protein G are the most enriched for detrimental mutations (1,080/1,585, 68.1%, OR=6.8, FET p<2.2×10^−16^), followed by proteins B (24/41, 58.6%, OR=2.7, FET p=1.2×10⁻^2^), A (1,318/1,926, 68.4%, OR=2.7, FET p<2.2×10^−16^), F (4,342/4,918, 88.2%, OR=1.9, FET p=1.6 × 10⁻^12^), and C (458/677, 67.7%, OR=1.4, FET p=4.8×10⁻^3^) (Fig. 4d). In Protein D, by contrast, nearly 90% (673/749) of interface mutations are detrimental, but there is no enrichment compared to non-interface surface residues (OR=1.5, FET p=5.6×10⁻²). Comparing different interfaces in the same protein, mutations in the F-G interface are more frequently detrimental than those in F-D interfaces (182/190, 95.8% vs 405/494, 82.0%; OR=5.0, FET p=6.2×10⁻⁷), likely due to additional roles of the F-G interaction beyond capsid assembly (*28*), whereas F-D interactions only provide transient scaffolding during procapsid assembly (*24*). In addition, mutations in residues involved in multiple interactions are more frequently detrimental than those involved in single interactions (1,746/1,969, 88.7% vs. 5,956/8,102, 73.5%; OR=2.8, FET p<2.2×10^−16^).

### Structural determinants of mutation tolerance

Using resolved and high-confidence predicted structures (see methods), protein cores and interaction interfaces together account for 73% of detrimental mutations across the ΦX174 proteome (Fig. 4a-b). A larger fraction of detrimental mutations maps to interaction interfaces than to protein cores (47.6% vs. 25.4%), but mutations in core residues are more likely to be detrimental than mutations in interfaces (5,058/5,783, 87.5% vs. 9,478/12,674, 74.8%; OR=2.4, FET p<2.2×10⁻¹⁶) (Fig. 4e), meaning that the greater importance of interface disruption is driven by the larger number of residues that participate in interfaces (43.2% of residues in interfaces vs. 18.3% of residues in cores). In ΦX174, therefore, ‘edgetic’ mutations (*36*) - those that disrupt molecular interactions rather than the folding of individual proteins - are the more frequent cause of detrimental mutations.

Across the proteome, proteins with a larger fraction of detrimental mutations in cores also show a larger fraction of detrimental mutations in interfaces (Spearman’s ρ=1.0, p=0.0167, n=5) (Fig. 4f). However, the relative contribution of core versus interface can vary substantially across proteins (Fig. 4a-c, g), indicating that detrimental effects are further shaped by specific protein functions. For example, the major spike protein G has the strongest enrichment of detrimental mutations at interfaces relative to non-interface surface residues (OR=6.8, FET p<2.2×10⁻¹⁶), but one of the weakest enrichments in core residues relative to surface residues (OR=2.3, FET p<2.2×10⁻¹⁶) (Fig. 4d). This pattern suggests that mutational sensitivity in G is shaped more by interactions than by core stability.

### Mutational effects are not well predicted by AI models

The complete mutagenesis of a viral proteome provides a large and well-calibrated dataset to evaluate the performance of computational methods for predicting variant effects. We evaluated variant effect predictors (VEPs) selected based on their state-of-the-art performance on viral proteins in ProteinGym (*37*). These include protein language models (PLMs), such as ESM-1v (*38*), Tranception-L (no retrieval) (*39*), VESPA (*40*), ProGen2 (*41*), ESM-2 (*42*), ESM3 (*43*), and ESM-C (*44*); evolutionary models based on multiple sequence alignments (MSAs), including GEMME (*45*), Tranception-L (with MSA retrieval) (*39*), PoET (200M) (*46*), and EVmutation2 (*47*); hybrid sequence-structure models, such as MIF-ST (*48*), SaProt (650M) (*49*) and ProSST (k=2048) (*50*); hybrid models integrating structure and MSAs, including ESCOTT (*51*), RSALOR (*52*), and ProteoCast (*53*); thermostability predictors, including DDMut (*54*), ThermoMPNN (*55*) and ProteinMPNN-ddG (*56*); an inverse-folding structure model, ESM-IF1 (*57*) and a binding affinity predictor, DDMut-PPI (*58*).

As a baseline predictor, we use per-residue RSA calculated from monomeric wildtype protein structures, a simple structural metric of mutational tolerance in protein domains (*32*). We also test RSA calculated on protein complexes, which captures interaction interfaces as solvent unaccessible. Predictor performance is evaluated using Spearman’s rank correlation coefficient (ρ) between predicted scores and experimentally measured fitness values for single-ORF amino acid substitutions in each ΦX174 protein (Fig. 4h). Because some predictors require structural inputs or features that are not available for all proteins, not every method can be evaluated across the full proteome (Fig. S4f).

Overall, the complex-derived RSA provides reasonable predictive performance (median ρ=0.41, n=7), outperforming the monomer-derived RSA baseline (median ρ=0.26, n=8). Among the evaluated VEPs, only two, ESCOTT (median ρ=0.50, n=4) and GEMME (median ρ=0.42, n=9), outperform the complex-derived RSA. ProteoCast (median ρ=0.41, n=7) that incorporates RSA into GEMME-derived scores shows comparable performance with the complex-derived RSA. RSALOR (median ρ=0.38, n=8), which also combines MSA-derived evolutionary information with RSA, performs better than the monomer-derived RSA, but worse than the complex-derived RSA. Protein stability predictors, including ThermoMPNN (median ρ=0.33, n=8), ProteinMPNN-ddG (median ρ=0.33, n=8), and DDMut (median ρ=0.30, n=8), also outperform the monomer-derived RSA baseline. Four PLM predictors show comparable performance with the monomer-derived RSA, including Progen2-xlarge (median ρ=0.34, n=9), VESPA-l (median ρ=0.32, n=9), Progen2-large (median ρ=0.31, n=10), and VESPA (median ρ=0.31, n=10).

Strikingly, therefore, nearly all state-of-the-art computational methods perform worse than the simple structure-based predictor of RSA calculated on protein complexes. This poor performance may, in part, reflect the underrepresentation of viral sequences in training datasets (*59*), the shallow sequence alignments of ΦX174, and the underrepresentation of ssDNA genomes in metagenomic data (*11*) (Fig. S4d-e).

### One quarter of damaging mutations have an unknown molecular mechanism

Structure predictions increase the fraction of detrimental mutations with a mechanistic interpretation from 54% to 73% (Fig. 4a-b), but a substantial fraction of detrimental mutations still lack a clear explanation (27%, 5,372/19,908). Of these, nearly half (47.8%; 2,570/5,372) are in protein regions with limited structural information (Fig. 4b, Table. S1), including those lacking high-confidence predicted structures, or missing from available resolved structures, such as the N-terminus of protein B and the C-terminus of protein E. The remaining 52.2% (2,802/5,372) occur at non-interface surface residues (Fig. 4b). These unexplained residues may include previously uncharacterized interfaces, regulatory regions, or other mechanisms not captured by current structural annotations. For example, some of the unexplained detrimental mutations in protein F (814 mutations) and protein A (1,305 mutations) may reflect hypothesized functional interactions of these proteins with the DNA replication machinery during genome packaging (*60*); and those in protein G (166 mutations) may relate to host lipopolysaccharide (LPS) recognition or interaction with protein H during infection initiation (*28*). In protein J, 161 detrimental mutations remain unexplained, 83.9% (135/161) of which occur at glycine residues. Glycine residues are also significantly enriched for detrimental mutations relative to non-glycine residues (135/152, 88.8% vs. 26/57, 45.6%, OR=9.5, FET p=3.5×10⁻^10^) in the non-interface surface, indicating that, in addition to capsid binding, conformational flexibility or backbone geometry is also important for protein J function (*61*).

Over 90% of unexplained mutations due to limited structural information are within the protein H (92.2%; 2,370/2,570). Protein H is the minor spike protein that oligomerizes into a decameric α-helical barrel that spans the cell membrane and mediates delivery of the viral DNA genome (*9*). However, the available resolved structure covers only ∼40% of the full-length protein (the coiled-coil region, aa 144–272) (*9*) and unexplained detrimental mutations (94.7%, 2,244/2,370) are mostly located in the structurally unresolved N- and C-terminal regions. The N-terminal region is enriched in hydrophobic residues reported to affect viral assembly and interactions of the protein with the lipid components of the host cell membrane (*62*) and detrimental mutations are enriched at hydrophobic residues throughout the protein (2,090/3,142, 66.5% vs. 1,479/3,032, 48.8%, OR=2.1, FET p<2.2×10⁻¹⁶) (Fig. S4g-h). Mutating inward-facing residues may also affect the interactions with DNA and interfere with genome delivery (*63*). Although these processes are likely dynamic and remain poorly characterized, our data provide a rich resource for generating mechanistic hypotheses and guiding future molecular modelling.

### Combining mutagenesis with structural modelling to understand protein A-DNA interactions

The replication of the ΦX174 genome has been extensively studied, serving as a model system for understanding rolling-circle DNA replication (*64*). However, the structures of the viral proteins A and C in the replication machinery have not been determined, leaving many mechanistic details unclear (*65*). We use our data in combination with AlphaFold 3 structure predictions (*66*) to further investigate these two proteins and their potential interactions during DNA replication.

Protein A is an HUH superfamily endonuclease that is characterised by conserved sequence motifs, including two histidine residues separated by a bulky hydrophobic residue and one or more catalytic tyrosine residues (*64*). It initiates rolling-circle replication by binding replicative form dsDNA and cleaving the viral (+) strand (Fig. 5a) (*67*). After each round of replication, protein A re-ligates the cleaved strand to release a circular ssDNA. Cleavage and religation occur at the nucleotide residues G4305 and A4306 within the 30nt origin of replication (ori^+^) (*68*, *69*) and require a divalent magnesium ion (Mg^2+^) (*64*). Two catalytic tyrosine residues, Tyr343 and Tyr347, mediate sequential nucleophilic attacks required for initiation, termination, and reinitiation of viral-strand processing (*70*). However, many mechanistic aspects beyond catalysis remain unclear, including how protein A engages dsDNA before cleavage, a process proposed to differ from its covalent ssDNA binding, in which a catalytic tyrosine forms a phosphotyrosine bond with the 5′ end of the cleaved viral strand (*71*).

Using AlphaFold 3 (*66*), we generated models for protein A bound to dsDNA (A-dsDNA) and ssDNA (A-ssDNA) (Fig. 5b-d). In the models with the highest A-DNA interface confidence scores, ranked by ipTM, protein A adopts a similar high-confidence fold (pTM=0.85 in the A-ssDNA model, and 0.86 in the A-dsDNA model), whereas its predicted ssDNA- and dsDNA-binding interfaces are distinct with slightly lower confidence (ipTM=0.83 in the A-ssDNA model, and 0.78 in the A-dsDNA model; Fig. S5a-b, h). While protein A circles around curved ssDNA in a C-clamped architecture, it binds dsDNA in a split-clamp shape, including disruption of base-pairing at the AT rich region (Fig. 5c-d). In both models, large DNA-interfaces are identified (81aa and 74aa respectively for dsDNA and ssDNA interfaces) with slightly more than half of the residues (44aa) participating in both interfaces.

The ssDNA-binding interface includes the previously identified catalytic residues Tyr343 and Tyr347 (*70*), as well as the HUH motif comprising His263 and His265, separated by Phe264 (Fig. 5c). The dsDNA-binding interface partially overlaps ssDNA-binding interface but does not include the HUH motif or the catalytic residues. Instead, dsDNA-binding interface includes residues that insert into the DNA duplex, potentially disrupting base paring and unwinding dsDNA (Fig. 5d). Detrimental mutations are enriched in the the shared residue interfaces (OR=1.92, p=4.1×10^−10^, n=442) and also in ssDNA-specific (OR=3.55, p<2.2×10⁻¹⁶, n=487) and dsDNA-specific interface residues (OR=1.76, p=1.7×10^−7^, n=405) (Fig. 5e). These results support the existence of distinct A-ssDNA and A-dsDNA interactions, both with functional importance.

The previously identified functional residues are highly sensitive to mutation. These include the catalytic residues Tyr343 and Tyr347 that mediate cleavage and religation of the viral strand (*70*), and the HUH motif residues His263 and His265, together with Glu254, which contact the Mg²⁺ near the catalytic site, potentially facilitating the reactions (Fig. 5c). Two juxtaposed threonine residues, Thr202 and Thr204, and a nearby Arg261 may further stabilize the DNA backbone or contribute to substrate positioning, possibly involving contacts between the threonine residues and thymine bases near the cleavage site (*69*, *72*).

The remaining DNA-interface, including the dsDNA-binding interface, is previously uncharacterized and contains multiple functional residues. The dsDNA-specific interface is centred on two juxtaposed phenylalanine residues Phe455 and Phe471, together with nearby positively charged residues, Lys454, Arg457, and Lys461, acting as a DNA strand-separation wedge (Fig. 5d). This organization resembles features of TATA-box binding protein (TBP), which binds the minor groove at the TATA box and induces DNA deformation through aromatic intercalation and backbone contacts (*73*, *74*). Similarly, in the A-dsDNA model, the juxtaposed phenylalanine residues may partially intercalate between base pairs and disrupt base stacking within the AT-rich region of ori^+^. Nearby positively charged residues, Lys454, Arg457 and Lys461, may stabilize the DNA backbone through electrostatic interactions. Together, these interactions promote duplex partially unwinding and induce a kinked DNA conformation resembling that observed in the A-ssDNA model (Fig. 5c).

### Protein C may allosterically regulate protein A activity

Protein C is an inhibitor of DNA replication initiation and is required for the transition from dsDNA replication to ssDNA genome synthesis and packaging (*30*). It has been suggested that protein C interacts with the initiation complex composed of Protein A, dsDNA, and the *E. coli* Rep helicase dimer (Fig. 5f) (*30*, *75*). The inhibition has been reported to require an intact Ori^+^ that is recognized and bound by protein A, although protein C has no effect on the DNA cleavage activity of protein A (*76*). More recently, cross-species complementation studies have suggested that protein C may directly interact with protein A (*65*). However, the mechanism by which protein C inhibits DNA replication, and whether this involves direct interaction with protein A or the broader initiation complex, remains unclear.

To address this, we focussed on potential A-C binding and generated a total of 150 models, 50 models for each of three increasingly complex assemblies by running AlphaFold 3 with 10 independent seeds per assembly: [1] the A-C complex; [2] the A-C-DNA complex; [3] the A-C-DNA-Rep complex, an initiation-complex-like assembly containing A, C, ssDNA, and *E. coli* Rep helicase dimers (Fig. S5e). The protein C structure is predicted with low confidence in isolation and remains low-confidence across all modelled complex structures (median pTM=0.53), unlike protein A which is predicted with high confidence in both A-C (median pTM=0.83) and A-C-DNA complexes (median pTM=0.84) (Fig. S5d, g). However, the inclusion of the Rep helicase dimer reduces protein A structure confidence (median pTM=0.6) (Fig. S5g). Consistent with this, protein A can adopt large structural variation in the initiation-complex-like assembly, including conformational changes (Fig. S5h). Despite the presence of Rep helicase, predicted A-C interfaces are retained in almost all the initiation-complex-like models (49/50). However, confidence in the predicted A-C interface is consistently low (median ipTM=0.18 for A-C alone, 0.34 for A-C-DNA, 0.17 for models including Rep proteins), and none reach the Alphafold confidence baseline (ipTM=0.6) (*66*).

To assess if any of the predicted complexes could provide plausible models for protein C function, we scored each model based on the enrichment of detrimental mutations at the predicted A-C interfaces in both proteins (Fig. S5f). Overall, 16 models have significant enrichment of detrimental mutations at the predicted A-C interface on protein A (Fig. S5i), and 33 have significant enrichment at the corresponding interface on protein C (FDR < 0.05) (Fig. S5j). Five of these models have enrichments for both proteins and all of them are models containing the Rep helicase. Interestingly, protein C is always positioned near the dsDNA-binding interface of protein A (Fig. 5h, Fig. S5k). The predicted A-C interactions are accompanied by a conformational change in protein A, in particular the residues implicated in DNA unwinding. Panel h in Figure 5 shows one of the supported models with the highest

A-C interface confidence score (ipTM=0.27). The structural module that protrudes into the DNA duplex and is predicted to promote local dsDNA unwinding is disrupted, whereas the catalytic sites remain preserved (Fig. 5h). Similar disruption or occlusion of the dsDNA-binding interface is observed in additional supported models, despite variation in protein A conformation and A-C interfaces, suggesting a potentially dynamic complex (Fig. S5k). Together, these mutagenesis-supported models suggest that protein C inhibits replication initiation by disrupting protein A-dsDNA engagement while leaving the catalytic site intact, allosterically regulating protein A function.

## Discussion

We have presented here a complete mutagenesis of a biological system, quantifying the fitness effects of all nucleotide and amino acid changes in the genome and proteome of the bacteriophage ΦX174. ΦX174 has served as a model system in molecular biology for more than 60 years and the complete mutagenesis of its genome and proteome adds to its historical importance as the first genome to be sequenced (*1*) and synthesised (*2*).

In total, more than half of nucleotide substitutions and over 60% of amino-acid substitutions impair viral fitness. The proportion of detrimental mutations varies widely over different genomic regions and proteins. Only a very small number of mutations increase fitness in the condition selected, suggesting this minimal three-replication-cycle selection assay closely matches the conditions to which the virus is adapted.

Two objectives of this study were to evaluate mechanistic causes of mutational effects in a well-studied system and how well they can be predicted by state-of-the-art machine learning models. Surprisingly, the whole proteome mutagenesis indicates that the most important mechanistic cause of fitness reduction is ‘edgetic’ mutations that disrupt interactions (*36*), with nearly 50% of all detrimental aa changes located in known and predicted interaction interfaces. This is almost double the number of detrimental mutations in individual protein cores, which account for ∼25% of damaging variants. At least in this system, disruption of interactions is more important than disruption of individual proteins as a cause of fitness loss. It will be interesting to re-evaluate the importance of interaction disruption in human genetic disease (*36*).

Despite the status of ΦX174 as a classic model system with extensive structural data, a quarter of detrimental protein mutations have no mechanistic explanation. Moreover, mutational effects across the proteome are only moderately-well predicted by current state-of-the art AI models, with most models outperformed by the simple structural baseline of relative solvent accessibility in protein complex structures. These rather humbling results are important, and further demonstrate the need for large-scale independent experimental evaluation of AI models.

By integrating mutagenesis with structural modelling, we provide mechanistic hypotheses for protein complexes involved in the initiation and inhibition of viral DNA synthesis. Our data suggest that protein A binds dsDNA and ssDNA differently: ssDNA-binding involves the catalytic site that mediates the cleavage and religation of viral strand, while dsDNA binding involves previous unidentified residues predicted to promote local DNA unwinding and formation of a kinked DNA conformation. Despite low computational confidence, mutational data support structural models for A-C interactions where protein C allosterically regulates protein A function, potentially as a dynamic complex. More generally, comprehensive mutagenesis could be used to evaluate the thousands of protein complex structure predictions that currently lack experimental support, guiding mechanistic interpretation, hypothesis generation, and downstream experimental design (*77–79*).

We believe that complete mutagenesis of biological systems has an important role to play in quantifying our current understanding of biology. Compared to other perturbation experiments, mutagenesis is cheap, highly-scalable, and generates quantitative parameter alterations in all biological polymers within a common experimental design. We have applied this approach to a landmark model virus, but one can envisage also applying it to complete pathways and regulatory networks (*80*). With highly scalable mutagenesis and editing technologies, it may also be applicable to entire prokaryotic (*81*) and eukaryotic (*82*) genomes.

## Methods

### Variant library design and preparation

The wild-type (WT) ΦX174 genome used in this study was derived from the commercial replicative-form I (RFI) ΦX174 genome (Promega, #D1531). This commercial genome carries an amber mutation in the lysis gene *E* (G587A) and differs from the canonical ΦX174 isolate Sanger reference genome (NC_001422.1) at four additional loci: relative to the reference, the Promega genome contains A833, G2731, T2811, and A3111, and only A3111 creates a non-synonymous amino-acid mutation in protein H (I61V). The amber mutation G587A was reverted using the Q5 Site-Directed Mutagenesis Kit (NEB, #E0554S) to restore the WT coding sequence. The corrected genome was validated by Oxford Nanopore sequencing and used as the reference backbone for all library construction. Genome annotations, including ORFs, promoters and terminators, were based on recent annotations reported in (*16*), except RBSs are based on (*1*).

For construction of the variant library pool, the genome was divided into 30 overlapping segments of 200-300 nt to design 30 mutant libraries (Fig. S1a-c). Collectively, these libraries contained all possible single nucleotide substitutions, as well as all possible 19 single-amino-acid substitutions and a stop codon for each codon. For each amino-acid substitution, the most frequently used *E. coli* codon was selected to minimize codon-usage bias (*83*). All synonymous codons encoding the wild-type amino acid at each position were also included.

Each of the 30 mutant libraries consisted of approximately 2,000 pooled mutant oligonucleotides synthesized as defined oligo pools by Twist Bioscience, with constant flanking regions for amplification and cloning (Fig. S1c). Oligo pools were PCR-amplified for 14 cycles using NEB Q5 Hot Start High-Fidelity DNA Polymerase (#M0493L), followed by ExoSAP (Thermo Fisher Scientific, #75001) treatment and purification using Qiagen MinElute Kit (#28006). For each library, three independent transformations were performed and followed by fitness selection as three independent biological replicates.

In parallel, 30 corresponding linearised ΦX174 WT backbone fragments were prepared for ligation to the mutant library. The linearised WT backbones were PCR-amplified using the a minimal amount of template DNA (0.5 ng) for 35 cycles using NEB Q5 Hot Start High-Fidelity DNA Polymerase (#E0554S), gel-purified using the Qiagen MinElute Gel Extraction Kit (#28606), and assembled with the matched mutant library pools using NEB Gibson Assembly Mix (#E2611L) according to the manufacturer’s instructions. Assembly products were then desalted using Millipore membrane filters (#VSWP02500), and 5ul from each library was electroporated into 50ul of home-made electrocompetent host bacterial strain *E.coli* C (ATCC #13706).

### High-throughput fitness selection

Selections began with the electroporation of assembled mutant genome libraries. Electroporated cells were immediately recovered in 1mL SOC medium (supplemented with 2.0 mM CaCl₂) at 37 °C for 40 min (corresponding to ∼1.5 generations). Recovered cultures were then mixed with 2 mL fresh log-phase host cells in TK media (5 g tryptone, 1.25 g NaCl, and 1.25 g KCl per 500 mL, supplemented with CaCl₂ to 2.0 mM) and incubated at 30°C for an additional 40 min to allow further phage infection and propagation. The total selection time was restricted to less than three wild-type generations (approximately 80 min) to minimize the possibility of compensatory mutations and spontaneous reversions. Mixed cultures were then centrifuged at 4 °C and 4,000 rpm for 20 min to pellet host cells, from which intracellular ΦX174 variant genomes were extracted as the selection output.

Transformation efficiency and library coverage were estimated by plating an aliquot of SOC-recovered culture using a double-layer TK agar method (15 mL bottom layer with 1.5% agar and 5 mL top layer with 0.7% agar). Plates were incubated at 30°C for 12-16 hours. Plaques, each representing an independent transformant, were counted by titration to estimate transformation efficiency and confirm adequate mutant coverage during selection.

### Next-generation Sequencing (NGS) library preparation

Intracellular ΦX174 variant genomes were extracted from the post-selection *E.coli* C pellets using the QIAGEN Miniprep Kit (#27106). For each library, matched input (desalted ligated product before electroporation) and output from the selection were quantified by quantitative PCR (qPCR) using Absolute SYBR Green Low ROX qPCR Mix (Thermo Fisher Scientific, #AB1323A). qPCR was performed with the same primer pairs used for oligo pool amplification, which annealed the constant flanking regions of each mutant library.

Normalised, equal amounts of DNA, corresponding to at least 20,000-fold coverage per variant, were used as templates for the first amplification step of NGS library preparation. This first PCR for NGS library preparation (PCR1) was performed for 15 cycles using library-specific primers containing Illumina adapters and frame-shift base overhangs by Q5 Hot Start High-Fidelity DNA Polymerase (#M0493L). PCR1 products were treated with ExoSAP (2 μL per 50 μL reaction) at 37 °C for 30 min, followed by heat inactivation at 80 °C for 20 min. Products were then purified using Qiagen MinElute Kit (#28006).

For the second PCR (PCR2), purified PCR1 products were amplified for 10 cycles at 62 °C using primers containing Illumina P5/P7 adapter sequences and unique sample indices by Q5 Hot Start High-Fidelity DNA Polymerase (#M0493L). PCR2 products were purified using the same protocol above, then quantified using a TapeStation D5000 assay, pooled equimolarly, gel-purified, and sequenced as 150-bp paired-end reads on an Illumina NovaSeq platform. A minimum median coverage of 1000 reads per variant was targeted.

### NGS data processing

Raw FASTQ files were processed using the DiMSum pipeline with default parameters (*84*). Designed variant lists were provided to restrict analysis to the intended variants using argument -barcodeIdentityPath. DiMSum quality-control diagnostics indicated that all single-nucleotide variant libraries passed the diagnostic error model.

### Fitness data normalization across libraries

After filtering out variants with fewer than 100 total input reads, fitness measurements and associated standard error for each variant were obtained through DiMSum output files (DiMSum_Project_fitness_replicates.RData). For each variant *i* in each mutation library *l*, the DiMSum-derived fitness and standard error were denoted as *F _i_*_,*l*_ and σ *_i_*_,*l*_ respectively.

A two-step normalization strategy was used to process these raw fitness values to place all measurements on a common scale. First, fitness values in each library were normalized using synonymous and nonsense variants as internally functional references.

Within-library normalization: internal nonsense/synonymous scaling (Eq. 1).

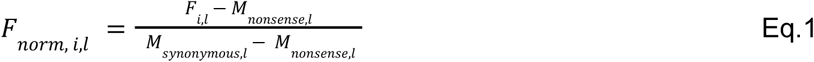

where *F _norm_*_, *i*,*l*_ is the normalized fitness of variant *i* in library *l*, *M _nonsense_*_,*l*_ is the mode of the nonsense fitness distribution in library *l*, and *M _synonymous_*_,*l*_ is the mode of the synonymous fitness distribution in library *l*.

The mode was defined as the fitness value corresponding to the maximum of the estimated density function. The synonymous and nonsense modes were estimated from their fitness distributions using kernel density estimation (KDE). This normalization step places the nonsense mode near 0 and the synonymous mode near 1 within each library.

Because this transformation is linear, the standard error of the mean fitness for each variant in each library derived from DiMSum pipeline (σ*_ij_*) was scaled by the same normalization factor (Eq.2).

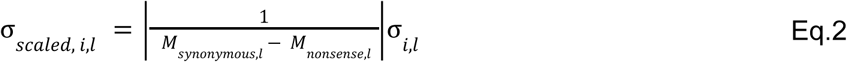

where

σ *_scaled_*_, *i*,*l*_ is the propagated error for variant *i* in library *l* after the first-stage within-library normalization.

Because essential and non-essential genes differ intrinsically in the fitness costs of nonsense variants, the initial within-library normalization, which corrects only for library-specific fitness range variation, could not establish a directly comparable scale across libraries. To bridge this gap, we used the mode of nonsense variants in single-ORF essential genes encoding structural proteins (i.e., proteins D, F, and G) as the library reference 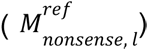 and rescaled the mean fitness (*F _scaled norm, i,l_*), as shown in Eq. (3).

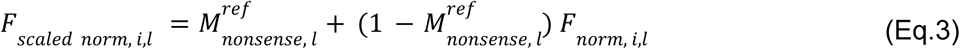

The corresponding standard error was rescaled again (σ *_rescaled_*_, *i*,*l*_) as in Eq. 4:

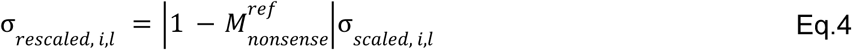

To align libraries with multimodal nonsense distributions or lacking a direct structural-protein reference, we implemented a sequential anchor-based rescaling procedure. The reference scale was first defined using structural proteins with similar nonsense-mode behavior (proteins F and G) and then propagated sequentially from G to H and from H to A using Eq. 3 and 4; for libraries exhibiting bimodal nonsense distributions, gene-specific nonsense modes were treated separately rather than pooled. Rescaling proceeded in order: starting with the G-only library, followed by the G+H library, then the A+H library (which lacks an essential structural reference), and finally the A-only library, whose A nonsense mode of the protein A from the libraries A+H and A+B was used as a reference to rescale. This generated a unified fitness dataset while preserving biologically meaningful differences across different proteins.

All scaled and normalised fitness effects for variants were combined into a single dataset (see ‘Data Availability’). For variants present in both libraries, the weighted mean 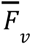 was computed using inverse-variance weights, and the variance 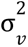 was propagated accordingly.

### Protein-level variant fitness effect calculation

For amino acid mutations with corresponding direct single-amino-acid variants, the measurements 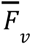 and 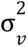 from the target single-ORF mutations were directly used.

For amino acid mutations with multiple genetic backgrounds, fitness effects of all the variants 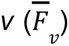 encoding the amino acid mutation *a* were averaged to obtain the mean fitness effects 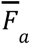, as shown in (Eq. 5).

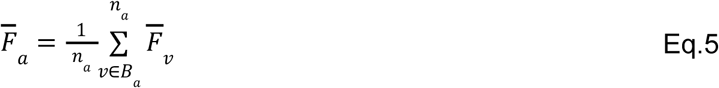

Here, *B_a_*denotes a given amino acid mutation and *n_a_* is the number of genetic variants encoding it, where a single nucleotide change can yield multiple amino acid substitutions across the phage proteome.

Standard error of mean for the fitness of each amino-acid substitution 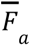, denoted as *SEM_a_*, is calculated by propagating the corresponding nucleotide-level errors 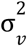 as shown in Eq. 6.

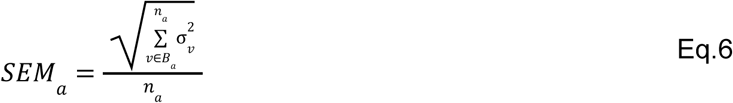

### Statistical Analysis

Statistical testing was performed independently for three datasets: all ΦX174 genomic variants, single-nucleotide (1nt) variants, and proteome-wide single-amino-acid (1aa) variants. For variants in each dataset, a two-sample student t-test was performed against a dataset-specific matching set of synonymous variants based on the mean fitness value and the corresponding errors.

For each dataset d, let *F_j,d_* denote the normalized fitness estimate of synonymous variant j, and let σ *_j_*_,*d*_ denote its propagated standard error. The synonymous reference fitness was estimated as an inverse-variance weighted mean:

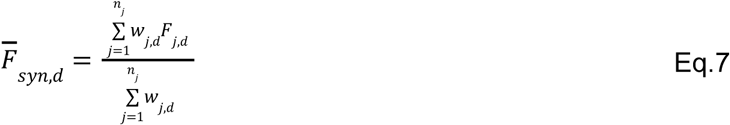

Where

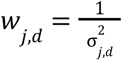

The propagated standard error of the synonymous reference was calculated as:

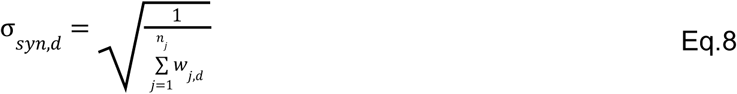

This produced three dataset-specific synonymous references correspondingly.

For each variant or mutation x in dataset d, the raw fitness effect relative to the synonymous reference was calculated as:

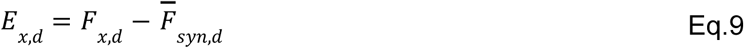

where *F _x_*_,*d*_ is the processed fitness value of variant or mutation x. Positive values indicate fitness above the synonymous reference, whereas negative values indicate fitness below the synonymous reference.

Statistical significance was assessed using a summary-statistic two-sample t-test comparing each variant or mutation with the corresponding dataset-specific synonymous reference:

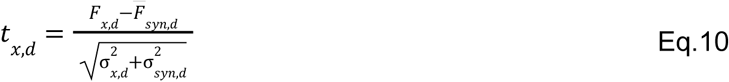

where σ *x*,*d* is the propagated measurement-level standard error of variant or mutation x, and σ *_syn_*_,*d*_ is the propagated standard error of the synonymous reference.

The degrees of freedom were approximated using the Welch-Satterthwaite equation:

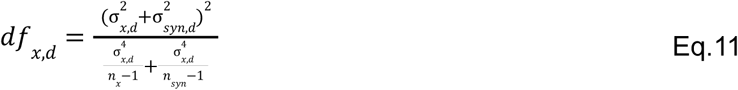

Because the propagated standard errors were derived from measurement-level uncertainty across three biological replicates, both sample-site terms *n_x_* and *n_sym_* were set to 3.

This choice is based on the degrees of freedom on the biological replicate structure of the experiment rather than on the number of collapsed genotypes, synonymous variants, or Multi-ORF backgrounds. This avoided inflating the degrees of freedom by treating aggregated observations as independent biological replicates.

Two-sided p-values were calculated from the t-distribution:

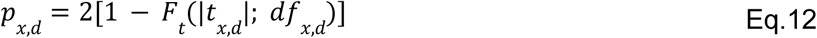

where *F_t_* is the cumulative distribution function of the t-distribution with *df _x_*_,*d*_ degrees of freedom.

Within each dataset, raw p-values were adjusted for multiple testing using the Benjamini-Hochberg false discovery rate (FDR). FDR correction was performed using the built-in R function ‘p.adjust’ with the method defined as ‘BH’. Thus, for each dataset d, the collection of raw p-values was adjusted to obtain FDR-adjusted p-values.

To calculate the statistical significance of enrichment of detrimental mutations in different protein structures, two-sided Fisher’s exact tests were performed on 2 × 2 contingency tables comparing detrimental versus non-detrimental mutations in each focal structural class against the corresponding reference class. Odds ratios were together calculated from the same contingency tables to quantify the direction and magnitude of enrichment.

### Mutation-effect classification

Variant effects were classified separately for each of the three datasets using fitness effects and FDR-adjusted significance. Variants with FDR ≥ 0.05 were classified as neutral. Among variants passing the FDR threshold, those with positive effects relative to the synonymous reference were classified as fitness-increasing, whereas those with negative effects were classified as fitness-reducing or detrimental. To further distinguish strong loss-of-function effects, statistically significant detrimental variants with a fitness decrease of at least 0.5 normalized fitness units relative to the synonymous reference were classified as lethal.

At the residue level, each nucleotide or amino acid position was classified as essential if at least one mutation at that position was lethal. Positions with no lethal mutations but at least one statistically significant detrimental mutation were classified as sensitive to mutation. Positions with complete mutational coverage and no statistically significant detrimental mutations were classified as tolerant. Positions with incomplete mutational coverage and no observed statistically significant detrimental mutations were left unclassified, because the effects of the missing mutations could not be inferred.

### Protein Structure analyses

Resolved structures of the ΦX174 procapsid including proteins B, D, F, and G (PDB: 1CD3) (*24*) and the mature virion capsid, including proteins F, G, and J (PDB: 2BPA) (*8*), were obtained from the Protein Data Bank. Additional structures included the coiled-coil region of protein H (residues 143-272; PDB: 4JPP) (*9*) and the “YES complex” containing a fragment of the lysis protein E (residues 1-65; PDB: 8G02) (*10*). Biological assembly files were used for all atomic distance calculations and interaction analyses. Structures containing multiple chains were separated by chain using ChimeraX and saved as individual PDB files for benchmarking of structure-based models. All molecular structures were visualized using ChimeraX (*85*).

Proteins without experimental structures, including proteins A (A*), C, K, as well as protein-protein and protein-DNA complexes, were predicted using the AlphaFold 3 web-server (https://alphafoldserver.com) (*66*) through the official webserver. For each, predictions were performed using 10 independent seeds, generating 5 ranked models per run. For protein monomers, the highest-ranked model with pTM > 0.8 was selected (only predictions for protein A monomer reach the standard), and models below this threshold were excluded.

Relative solvent accessibility (RSA) was calculated using PyMOL API per-residue surface area computation action. Residues with RSA < 25% were classified as core, whereas residues with RSA ≥ 25% were classified as surface. Surface residues were further classified as interfaces if any two heavy atoms of interacting proteins or DNA molecules are within 5 Å distance. This distance-based contact analysis is performed using ChimeraX, with the built-in contact analysis (*85*). Remaining surface residues were classified as non-interacting surfaces.

### Structure modelling using AlphaFold 3

Protein-DNA and protein-protein complex structures were modelled using the AlphaFold 3 server (https://alphafoldserver.com) (*66*). To model protein A bound to ssDNA (A-ssDNA), we used the wild-type protein A sequence together with a 30-nt ori^+^ sequence corresponding to the origin of replication (ori^+^) for the viral (+) strand. To model protein A bound to dsDNA (A-dsDNA), we used the wild-type protein A sequence together with the same 30-nt ori^+^ sequence and its reverse-complementary sequence as a paired DNA duplex. All protein A-DNA predictions included an Mg^2+^ ion. For each interaction, predictions were again performed using 10 independent seeds, with five internally ranked models generated per seed. The protein A structure is predicted with high confidence in both A-dsDNA and A-ssDNA complexes (median predicted TM-scores (pTM) = 0.86 for A-dsDNA models and 0.84 for A-ssDNA models) (Fig. S5a-b). However, confidence scores in the predicted protein-DNA interfaces are lower and more variable (Fig. S5a). Across both model sets, predicted A-DNA interfaces are well supported by enrichment of detrimental mutations (Fig. S5c). The A-ssDNA and A-dsDNA structures presented in the main text corresponded to model0 generated with seed5 and model0 generated with seed2, respectively.

Protein A-C interactions were modelled in three functional contexts (Fig. S5e-f). First, A-C complexes were modelled using the wild-type sequences of proteins A and C. Second, A-C-DNA complexes were modelled by adding the 30-nt ori^+^ DNA sequence. Third, initiation-complex-like assemblies were modelled by including an *E. coli* Rep helicase dimer (*75*). All proteins A-C complex predictions included an Mg²⁺ ion to be consistent with the protein A-DNA modelling conditions and to allow comparison across predicted complexes. For each context, predictions were performed using 10 independent seeds, with five ranked models generated per seed. When calculating the enrichment of detrimental mutations across each structural class of protein A in the A-C complexes, residues involved in A-DNA interfaces were excluded. The five selected A-C interaction structures presented in the main text corresponded to model3 generated with seed8, models0 and 4 generated with seed9, model3 generated with seed2, and model2 generated with seed4 (see ‘Data Availability’).

### Multiple sequence alignments (MSAs) for ΦX174 proteins

Multiple sequence alignments (MSAs) for ΦX174 proteins were generated using JackHMMER as implemented in the EVcouplings pipeline (*86*). For each ΦX174 protein, the wild-type amino-acid sequence was submitted to the EVcouplings webserver (https://v2.evcouplings.org). Separate searches were performed against the UniRef100, UniRef90, and MGnify+UniProt databases using the default webserver settings, except that the bitscore threshold was varied from 0.1 to 0.9 in increments of 0.1. Each search was run for five JackHMMER iterations.

One MSA per protein was selected using one fixed rule: among alignments with an effective depth per residue greater than one (defined as *Meff*/*L* > 1), we chose the highest bitscore threshold, favouring stringency for closer homologs while preserving depth; if none met this cutoff, we selected the alignment with the largest available *Meff*/*L*.The effective number of sequences was calculated by EVcouplings using a sequence-identity threshold of θ = 0. 8. The selected MSA files are summarized in Supplementary Table 2 and were only used as retrieval inputs for benchmarking Tranception with MSA retrieval (see below).

### Evaluation of variant effect predictors (VEP)

Variant effect predictors were run using wild-type ΦX174 protein sequences, single-ORF amino acid mutation lists, and, where required, experimentally resolved or high-confidence predicted protein structures as inputs. Protein K was excluded from benchmarking because no fitness-defective missense variants were detected in our assay. Unless otherwise stated, predictors were run with their pretrained weights and default inference settings.

Protein language models were run using wild-type ΦX174 protein sequences as input. VESPA was run following the official GitHub instructions and generated both VESPA and VESPA-l scores for each mutation. ESM-1v was evaluated in both single-model and ensemble modes. The first ESM-1v model, esm1v_t33_650M_UR90S_1, was used for the single-model score, whereas the ensemble score was calculated by averaging scores across all five ESM-1v models. ESM-2 was evaluated using the 8M, 35M, 150M, 650M, 3B, and 15B parameter models. ESM3-open was evaluated using the 1.4B parameter model, and ESM-Cambrian was evaluated using the 300M, 600M, and 6B parameter models. Progen2 was evaluated using the large, 2.7B, and xlarge, 6.4B, models, generating Progen2-L and Progen2-XL scores, respectively. These protein language models were run in Google Colab.

Evolutionary models based on multiple sequence alignments (MSAs) were run using MSA files generated by the corresponding predictor workflows or webservers, unless otherwise specified. GEMME was run through the official GEMME webserver by submitting each ΦX174 protein sequence (https://www.lcqb.upmc.fr/GEMME). GEMME-generated MSA files were reused as inputs for ProteoCast (https://proteocast.ijm.fr/) and ESCOTT (https://prescott.lcqb.upmc.fr). PoET was evaluated using the 200M-parameter model implemented in OpenProtein (https://www.openprotein.ai/). EVmutation2 was evaluated using evedesign (https://evedesign.bio/). Tranception Large (Tranception-L), a 700M parameter model, was evaluated with and without MSA retrieval in Google Colab. Tranception was the only predictor run with manually selected MSA inputs for retrieval mode, using the MSA selection criteria described above.

Structure-aware predictors were evaluated using experimentally resolved or high-confidence predicted structures as inputs. MIF-ST, ESM-IF1, and ProSST (k=2048) were run using the Directed Evolution module in the VenusFactory2 (https://venusfactory.cn/playground). SaProt was evaluated using the 650M-parameter SaProt in the official Google Colab notebook. ProteoCast was evaluated through its webserver (https://proteocast.ijm.fr/) using GEMME-derived MSA files and structure files as inputs. ESCOTT was evaluated through its official webserver using GEMME-derived MSA files and structure files as inputs. RSALOR was run using the official Google Colab notebook. DDMut was evaluated through the official DDMut webserver by submitting mutation-list files and corresponding protein structures (https://biosig.lab.uq.edu.au/ddmut/). DDMut-PPI was run through the official DDMut-PPI webserver for protein complexes with available structural inputs (https://biosig.lab.uq.edu.au/ddmut_ppi/). ThermoMPNN was run using the official Colab notebook with protein structures as input. ProteinMPNN-ddG was evaluated through Neurosnap (https://neurosnap.ai/) by submitting protein structures and mutation lists.

As a technical control, we included blaTEM-1 β-lactamase variant effects (*87*), which is also included in ProteinGym. For each predictor, blaTEM-1 scores were compared with the corresponding published or ProteinGym benchmark scores (*37*). The positive correlations and near-identical score distributions obtained for blaTEM-1 served as a control that the predictors were being run correctly.

## Supplementary figure legends

**Figure S1:**
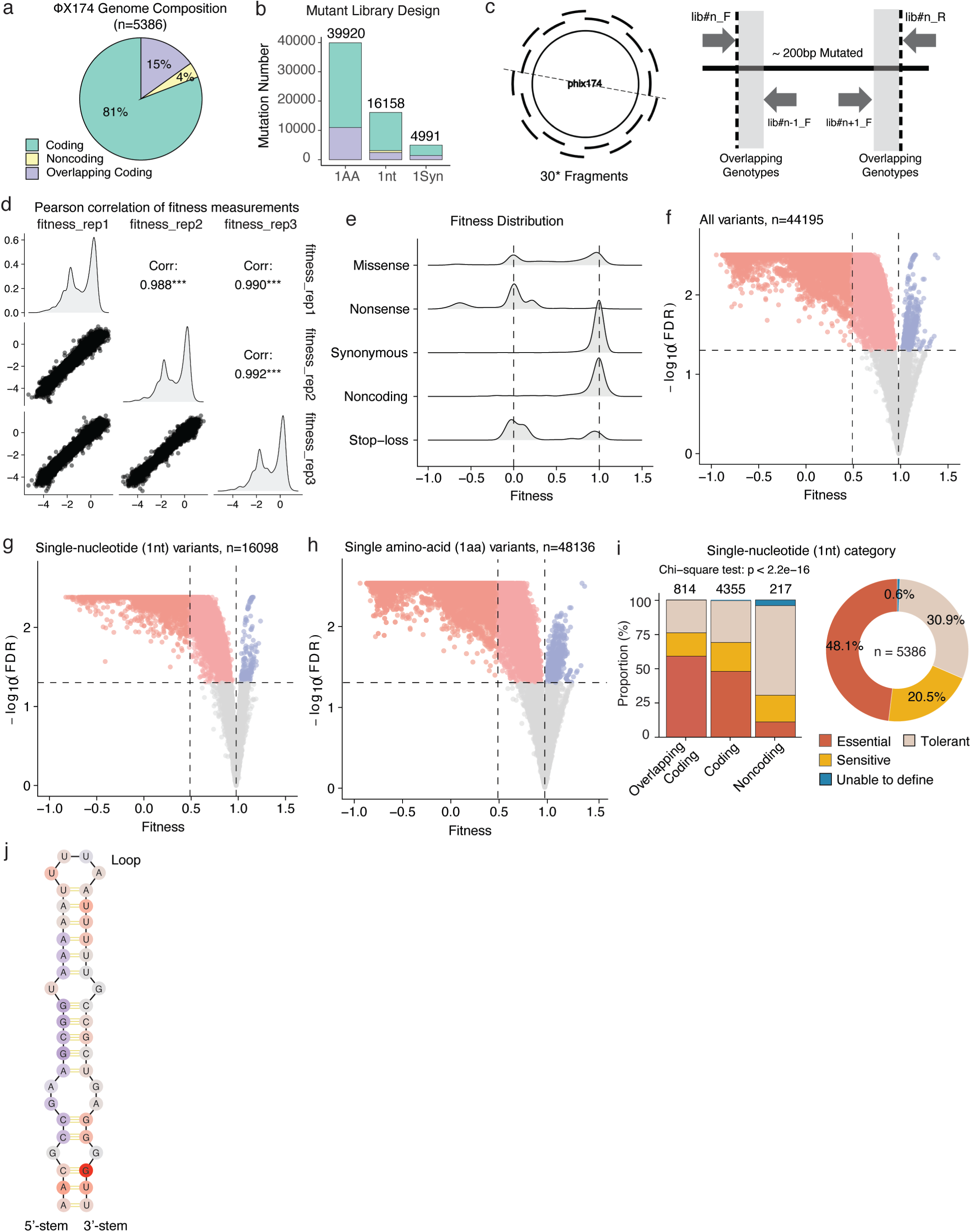
Overview of mutant library design and data analysis. a, ΦX174 genome composition. b, Composition of the mutant library. c, Mutant library design and construction strategy for mutating the circular ΦX174 genome. d, Replicate correlations. e. Distribution of variant fitness effects at the ΦX174 variant level. f-h, Volcano plots showing mutation-effect classification based on fitness values and FDR-adjusted significance at the ΦX174 variant level (f), single-nucleotide variant level (g), and single-amino-acid variant level (h). i, Nucleotide residue category. j, RNAfold-predicted hairpin structure of the TF transcriptional terminator (*88*). Each circle represents a nucleotide residue and is colored by the lowest fitness effect observed among mutations at that position.

**Figure S2:**
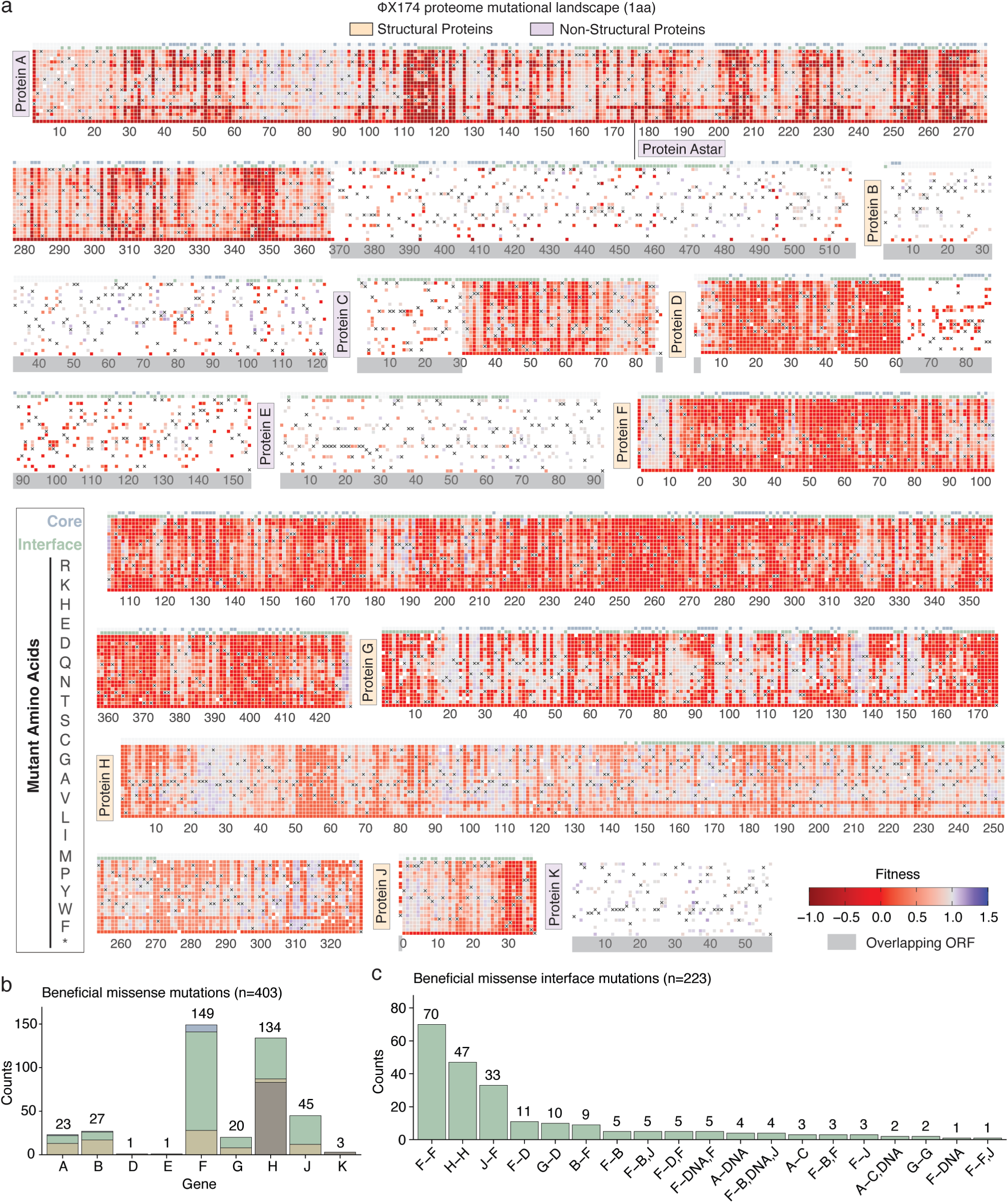
Amino-acid variant effects and beneficial variants. a, Heatmaps showing amino-acid variant fitness landscapes for 11 viral proteins, with multi-ORF variants removed. b, Distribution of beneficial amino-acid variants across viral proteins. c, Distribution of beneficial amino-acid variants across protein interfaces.

**Figure S3:**
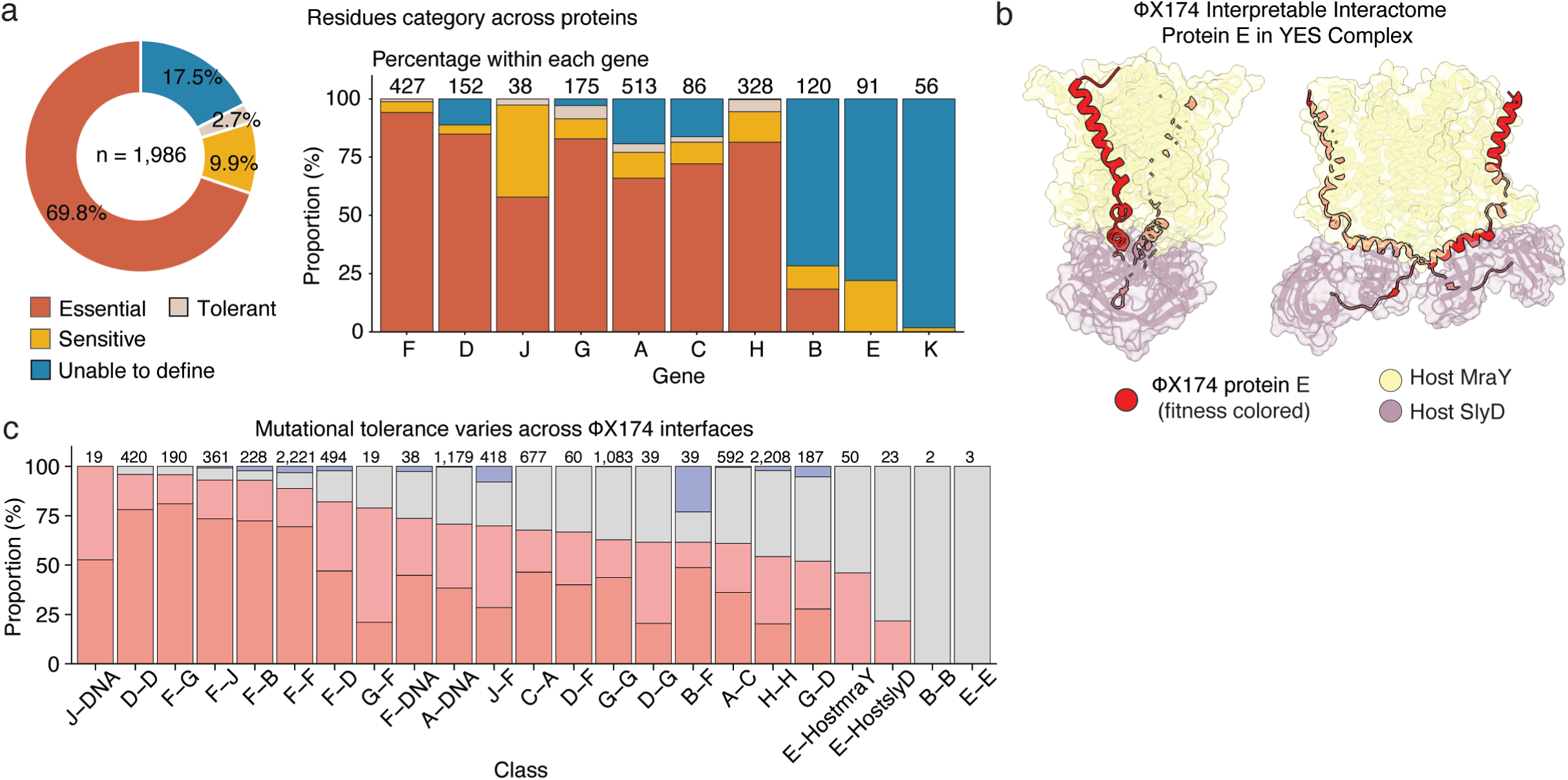
Amino-acid residue classification and overview of protein-interface variants. a, Amino-acid residue categories. b, Mutation categories for single amino-acid substitutions (single-ORF, missense mutations) across protein interfaces. c, Resolved structure for ΦX174 protein E in the ‘YES’ complex from PDB: 8G02 (*10*), and protein E is colored by positional median fitness effect.

**Figure S4:**
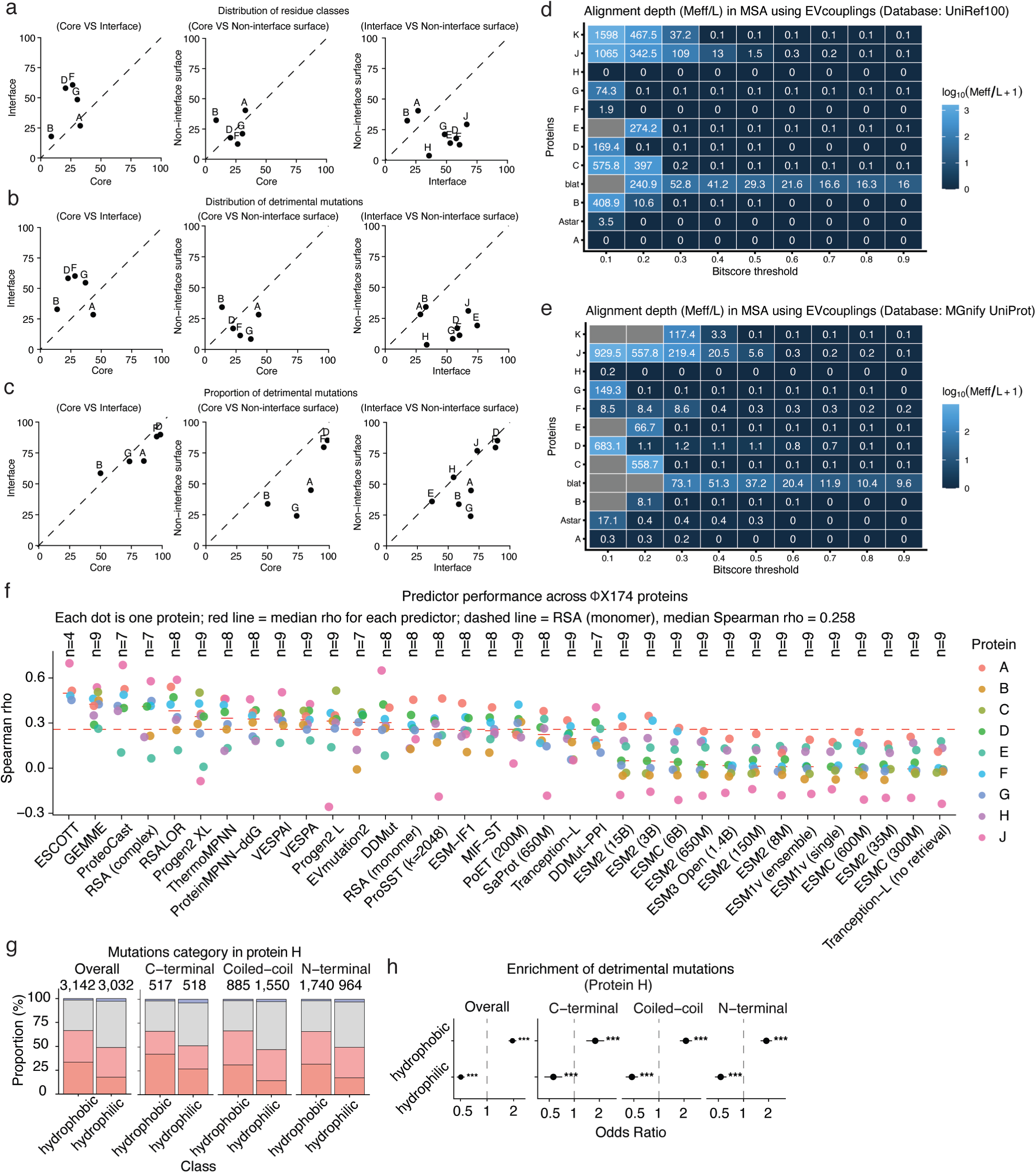
Mechanistic interpretation of detrimental amino-acid variants and evaluation of variant effect predictors. a, Proportion of core and interface residues across viral proteins. b, Relationships between the distribution of detrimental mutations in core vs. interface, core vs. non-interface surface, and interface vs. non-interface surface. c, Relationships between the proportions of detrimental mutations in core vs. interface, core vs. non-interface surface, and interface vs. non-interface surface. d-e, Multiple sequence alignments generated by EVcoulings using UniRef100 (d) and MGnify plus UniProt (e). f, Evaluation of variant effect predictors. Predictors are ordered from left to right by their median performance across available proteins. Each point represents one protein and is colored by protein identity. g, Mutation categories for single amino-acid substitutions (single-ORF, missense mutations) in protein H. h, Enrichment of detrimental mutations in hydrophobic vs. hydrophilic residues in protein H.

**Figure S5:**
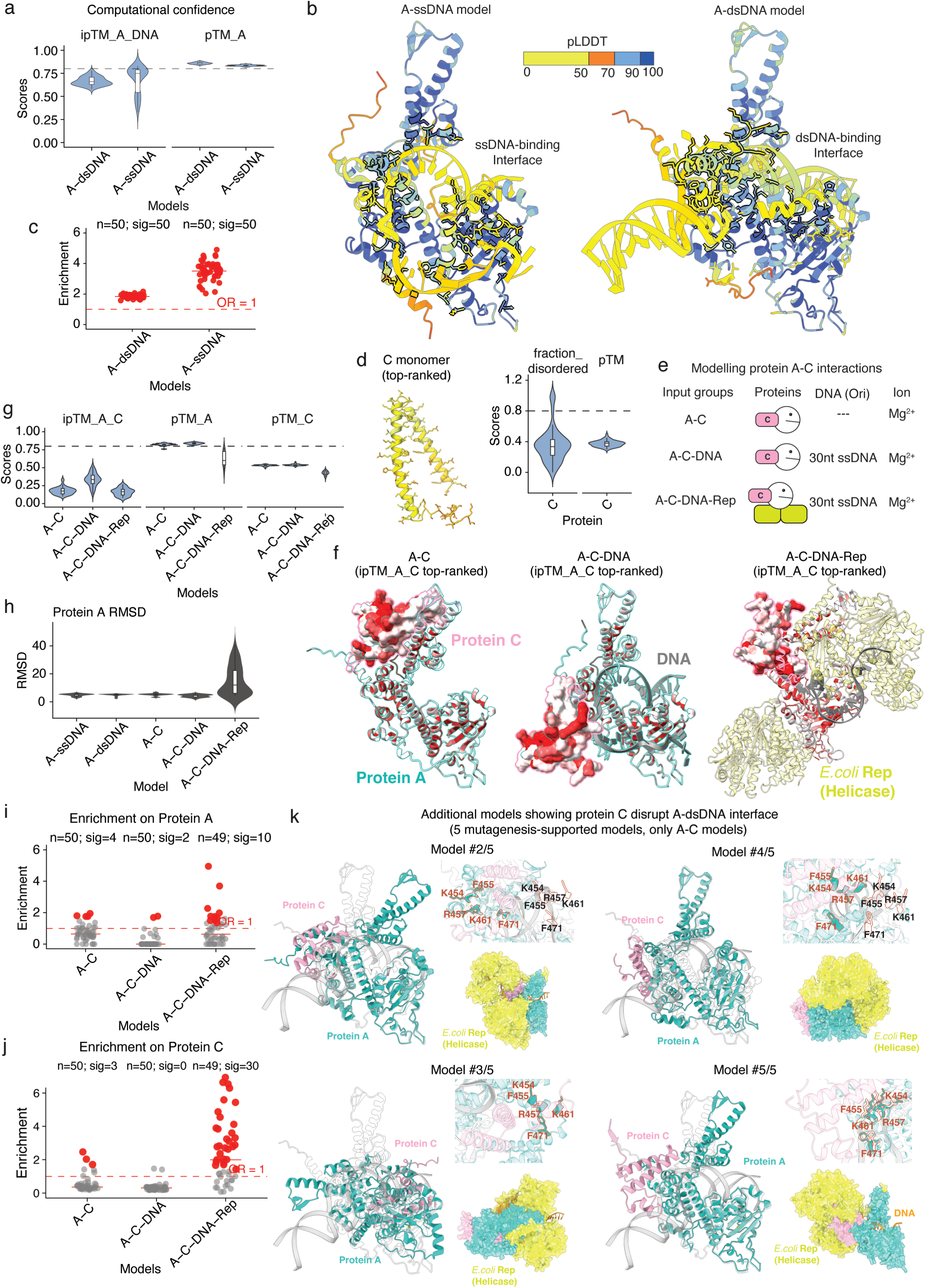
Evaluating and scoring structure predictions using comprehensive mutagenesis. a, AlphaFold 3 confidence metrics for A-ssDNA and A-dsDNA models, including predicted TM-scores (pTM) and interface predicted TM-scores (ipTM). b, Selected A-ssDNA and A-dsDNA models colored by predicted local distance difference test (pLDDT). c, Enrichment of detrimental mutations in predicted A-DNA interfaces in A-ssDNA and A-dsDNA models. d, AlphaFold 3 modelling of monomeric protein C. e, AlphaFold 3 modelling of A-C interactions in the three assembly contexts. f, Example structures from each A-C modelling context, with proteins A and C colored by positional median fitness effect. g. AlphaFold 3 confidence metrics for A-C complex models. h. Structural alignment of protein A across models, using the selected A-ssDNA model as the reference structure; alignments are summarized by RMSD calculated across all 513 amino-acid residues of protein A. i-j, Enrichment of detrimental mutations in predicted A-C interfaces on protein A (i) and protein C (j). k, Additional mutagenesis-supported structure models for A-C interactions.

---

**Supplementary Table 1.**

| Protein | Protein category | Primary Functions | Structure | Coverage |
| --- | --- | --- | --- | --- |
| A | Replication<br>(Non-structural) | Endonuclease, initiates and terminates the rolling-circle DNA replication | AlphaFold 3<br>(pTM > 0.8) | 100%<br>(513/513) |
| A* | Replication<br>(Non-structural) | Inhibits host cell DNA replication | - | - |
| B | Internal Scaffolding<br>(Structural) | Supports procapsid assembly | PDB: 1CD3<br>(residues: 1-8,61-120) | 56.7%<br>(68/120) |
| C | DNA Replication<br>(Non-structural) | Inhibits replication initiation and regulates the switch from dsDNA replication to genomic ssDNA synthesis | AlphaFold 3<br>(only potential interfaces) | 65.1%<br>(56/86) |
| D | External Scaffolding<br>(Structural) | Mediates procapsid assembly and forms a temporary outer shell | PDB: 1CD3<br>(residues: 5-152) | 97.4%<br>(148/152) |
| E | Host cell lysis<br>(Non-structural) | Inhibits bacterial cell wall synthesis | PDB: 8G02<br>(residues: 1-65) | 70.3%<br>(64/91) |
| F | Major capsid<br>(Structural) | Forms the main icosahedral shell protecting the viral genome | PDB: 1CD3<br>PDB: 2BPA<br>(residues: 2-427) | 99.8%<br>(426/427) |
| G | Major spike<br>(Structural) | Forms pentameric surface spikes that recognize and bind the host cell | PDB: 1CD3<br>PDB: 2BPA | 100%<br>(175/175) |
| H | Minor spike<br>(Structural) | Forms a decameric molecular conduit to inject the viral DNA into the host cell | PDB: 4JPP<br>(residues 144-272) | 38.3%<br>(129/328) |
| J | DNA-binding<br>(Structural) | Binds and guides the circular ssDNA genome inside the capsid during packaging | PDB: 2BPA<br>(residues: 3-38) | 94.7%<br>(36/38) |
| K | Burst-size control<br>(Non-structural) | Enhances burst size, though not strictly required for infection | No structure | 0%<br>(0/56) |

**Supplementary Table 2.**

| Protein | Meff/L | Bitscores | Searching Database |
| --- | --- | --- | --- |
| A | 0.3 | 0.2 | MGnify+UniProt |
| A* | 17.1 | 0.1 | MGnify+UniProt |
| B | 8.1 | 0.2 | MGnify+UniProt |
| C | 558.7 | 0.2 | MGnify+UniProt |
| D | 1.1 | 0.5 | MGnify+UniProt |
| E | 66.7 | 0.2 | MGnify+UniProt |
| F | 8.6 | 0.3 | MGnify+UniProt |
| G | 149.3 | 0.1 | MGnify+UniProt |
| H | 0.2 | 0.1 | MGnify+UniProt |
| J | 5.6 | 0.5 | MGnify+UniProt |
| K | 3.3 | 0.4 | MGnify+UniProt |

## Supplementary Materials

**File 1.** Wild-type (WT) ΦX174 genome sequence used in this study.

**File 2.** ΦX174 variant annotations and mutant library metadata.

**File 3.** Fitness measurements for all ΦX174 variants.

**File 4.** Fitness measurements for single-nucleotide (1nt) variants.

**File 5.** Residue-level summary of single-nucleotide (1nt) mutational effects.

**File 6.** Fitness measurements for single-amino-acid (1aa) variants.

**File 7.** Residue-level summary of single-amino-acid (1aa) mutational effects.

**File 8.** Performance evaluation of variant-effect predictors.

**File 9.** Variant-effect predictions generated using sequence- and sequence-alignment-based methods.

**File 10.** Variant-effect predictions generated using structure-based methods.

**File 11.** Selected AlphaFold 3 structural models of ΦX174 protein A-ssDNA, A-dsDNA, and proteins A-C interaction complexes.

## Acknowledgements

This work was supported by Wellcome (Grant reference: 220540/Z/20/A,‘Wellcome Sanger Institute Quinquennial Review 2021-2026’), a European Research Council (ERC) Advanced (883742) grant, the Spanish Ministry of Science and Innovation (LCF/PR/HR21/52410004, EMBL Partnership, Severo Ochoa Center of Excellence), Agència de Gestió d’Ajuts Universitaris i de Recerca (AGAUR, 2021 SGR01226), and the CERCA Program/Generalitat de Catalunya. H.W. was supported by a FPI fellowship PREP2023-001380, financed by MICIU/AEI /10.13039/501100011033 y por el FSE+.

## Author contributions

H.W. performed all experiments and analyses. H.W., X.L. and B.L. designed analyses and wrote the paper. X.L. and B.L. conceived and supervised the project.

## Data availability

All sequencing data have been deposited in the European Nucleotide Archive (ENA) at EMBL-EBI under accession number PRJEB120835.

## Code availability

All scripts used in this study are available at: https://github.com/lehner-lab/phix174_WGM.

## References

1. F. Sanger, G. M. Air, B. G. Barrell, N. L. Brown, A. R. Coulson, Nucleotide sequence of bacteriophage ϕX174 DNA. Nature (1977).

2. H. O. Smith, C. A. Hutchison 3rd, C. Pfannkoch, J. C. Venter, Generating a synthetic genome by whole genome assembly: phiX174 bacteriophage from synthetic oligonucleotides. Proc. Natl. Acad. Sci. U. S. A. 100, 15440–15445 (2003).

3. V. Sertic, N. Boulgakov, Classification et identification des typhi-phages. CR Soc. Biol. Paris 119, 1270–1272 (1935).

4. R. L. Sinsheimer, A single-stranded deoxyribonucleic acid from bacteriophage φX174. J. Mol. Biol. 1, 43–IN6 (1959).

5. B. G. Barrell, G. M. Air, C. A. Hutchison 3rd, Overlapping genes in bacteriophage phiX174. Nature 264, 34–41 (1976).

6. S. M. Doore, B. A. Fane, The microviridae: Diversity, assembly, and experimental evolution. Virology 491, 45–55 (2016).

7. T. Dokland, R. McKenna, L. L. Ilag, B. R. Bowman, N. L. Incardona, B. A. Fane, M. G. Rossmann, Structure of a viral procapsid with molecular scaffolding. Nature 389, 308–313 (1997).

8. R. McKenna, D. Xia, P. Willingmann, L. L. Ilag, S. Krishnaswamy, M. G. Rossmann, N. H. Olson, T. S. Baker, N. L. Incardona, Atomic structure of single-stranded DNA bacteriophage phi X174 and its functional implications. Nature 355, 137–143 (1992).

9. L. Sun, L. N. Young, X. Zhang, S. P. Boudko, A. Fokine, E. Zbornik, A. P. Roznowski, I. J. Molineux, M. G. Rossmann, B. A. Fane, Icosahedral bacteriophage ΦX174 forms a tail for DNA transport during infection. Nature 505, 432–435 (2014).

10. A. K. Orta, N. Riera, Y. E. Li, S. Tanaka, H. G. Yun, L. Klaic, W. M. Clemons Jr, The mechanism of the phage-encoded protein antibiotic from ΦX174. Science 381, eadg9091 (2023).

11. P. C. Kirchberger, H. Ochman, Microviruses: A world beyond phiX174. Annu. Rev. Virol. 10, 99–118 (2023).

12. P. R. Jaschke, E. K. Lieberman, J. Rodriguez, A. Sierra, D. Endy, A fully decompressed synthetic bacteriophage øX174 genome assembled and archived in yeast. Virology 434, 278–284 (2012).

13. M. S. Faber, J. T. Van Leuven, M. M. Ederer, Y. Sapozhnikov, Z. L. Wilson, H. A. Wichman, T. A. Whitehead, C. R. Miller, Saturation Mutagenesis genome engineering of infective ΦX174 bacteriophage via unamplified oligo pools and Golden Gate assembly. ACS Synth. Biol. 9, 125–131 (2020).

14. B. Álvarez-Rodríguez, S. Velandia-Álvarez, C. Toft, R. Geller, Mapping mutational fitness effects across the coxsackievirus B3 proteome reveals distinct profiles of mutation tolerability. PLoS Biol. 22, e3002709 (2024).

15. W. Bakhache, W. Symonds-Orr, L. McCormick, P. T. Dolan, Deep mutation, insertion and deletion scanning across the Enterovirus A proteome reveals constraints shaping viral evolution. Nat. Microbiol. 10, 158–168 (2025).

16. D. Y. Logel, P. R. Jaschke, A high-resolution map of bacteriophage øX174 transcription. Virology 547, 47–56 (2020).

17. K. Arai, A. Kornberg, Unique primed start of phage phi X174 DNA replication and mobility of the primosome in a direction opposite chain synthesis. Proc. Natl. Acad. Sci. U. S. A. 78, 69–73 (1981).

18. C. B. Harley, R. P. Reynolds, Analysis of E. coli promoter sequences. Nucleic Acids Res. 15, 2343–2361 (1987).

19. E. A. Campbell, O. Muzzin, M. Chlenov, J. L. Sun, C. A. Olson, O. Weinman, M. L. Trester-Zedlitz, S. A. Darst, Structure of the bacterial RNA polymerase promoter specificity sigma subunit. Mol. Cell 9, 527–539 (2002).

20. M. I. Voskuil, G. H. Chambliss, The TRTGn motif stabilizes the transcription initiation open complex. J. Mol. Biol. 322, 521–532 (2002).

21. Y.-H. Huang, C.-Y. Huang, Structural insight into the DNA-binding mode of the primosomal proteins PriA, PriB, and DnaT. Biomed Res. Int. 2014, 195162 (2014).

22. G. S. Goetz, T. Schmidt-Glenewinkel, M. H. Hu, N. Belgado, J. Hurwitz, Studies on the role of the phi X174 gene A protein in phi X viral strand synthesis. II. Effects of DNA replication of mutations in the 30-nucleotide icosahedral bacteriophage origin. J. Biol. Chem. 263, 16433–16442 (1988).

23. Y. C. Shin, G. F. Bischof, W. A. Lauer, R. C. Desrosiers, Importance of codon usage for the temporal regulation of viral gene expression. Proc. Natl. Acad. Sci. U. S. A. 112, 14030–14035 (2015).

24. T. Dokland, R. A. Bernal, A. Burch, S. Pletnev, B. A. Fane, M. G. Rossmann, The role of scaffolding proteins in the assembly of the small, single-stranded DNA virus phiX174. J. Mol. Biol. 288, 595–608 (1999).

25. D. R. Rokyta, C. L. Burch, S. B. Caudle, H. A. Wichman, Horizontal gene transfer and the evolution of microvirid coliphage genomes. J. Bacteriol. 188, 1134–1142 (2006).

26. J. T. Van Leuven, J. S. Patel, C. Beard, F. M. Ytreberg, L. Scott, K. Burns, E. Altman, Y. Sapozhnikov, C. F. Tovissodé, J. Yang, H. A. Wichman, B. M. Rubenstein, C. R. Miller, ΦX174 bacteriophage viability predicted by protein biophysical modeling, bioRxivorg (2025). 10.64898/2025.12.01.691700.

27. D. Rennell, S. E. Bouvier, L. W. Hardy, A. R. Poteete, Systematic mutation of bacteriophage T4 lysozyme. J. Mol. Biol. 222, 67–88 (1991).

28. Y. Sun, A. P. Roznowski, J. M. Tokuda, T. Klose, A. Mauney, L. Pollack, B. A. Fane, M. G. Rossmann, Structural changes of tailless bacteriophage ΦX174 during penetration of bacterial cell walls. Proc. Natl. Acad. Sci. U. S. A. 114, 13708–13713 (2017).

29. S. Eisenberg, A. Kornberg, Purification and characterization of phiX174 gene A protein. A multifunctional enzyme of duplex DNA replication. J. Biol. Chem. 254, 5328–5332 (1979).

30. A. Aoyama, M. Hayashi, Synthesis of bacteriophage phi X174 in vitro: mechanism of switch from DNA replication to DNA packaging. Cell 47, 99–106 (1986).

31. S. Gillam, T. Atkinson, A. Markham, M. Smith, Gene K of bacteriophage phi X174 codes for a protein which affects the burst size of phage production. J. Virol. 53, 708–709 (1985).

32. A. Beltran, X. ’er Jiang, Y. Shen, B. Lehner, Site-saturation mutagenesis of 500 human protein domains. Nature 637, 885–894 (2025).

33. K. Tsuboyama, J. Dauparas, J. Chen, E. Laine, Y. Mohseni Behbahani, J. J. Weinstein, N. M. Mangan, S. Ovchinnikov, G. J. Rocklin, Mega-scale experimental analysis of protein folding stability in biology and design. Nature 620, 434–444 (2023).

34. A. David, M. J. E. Sternberg, The contribution of missense mutations in core and rim residues of protein-protein interfaces to human disease. J. Mol. Biol. 427, 2886–2898 (2015).

35. A. J. Faure, J. Domingo, J. M. Schmiedel, C. Hidalgo-Carcedo, G. Diss, B. Lehner, Mapping the energetic and allosteric landscapes of protein binding domains. Nature 604, 175–183 (2022).

36. Q. Zhong, N. Simonis, Q.-R. Li, B. Charloteaux, F. Heuze, N. Klitgord, S. Tam, H. Yu, K. Venkatesan, D. Mou, V. Swearingen, M. A. Yildirim, H. Yan, A. Dricot, D. Szeto, C. Lin, T. Hao, C. Fan, S. Milstein, D. Dupuy, R. Brasseur, D. E. Hill, M. E. Cusick, M. Vidal, Edgetic perturbation models of human inherited disorders. Mol. Syst. Biol. 5, 321 (2009).

37. P. Notin, A. W. Kollasch, D. Ritter, L. van Niekerk, S. Paul, H. Spinner, N. Rollins, A. Shaw, R. Weitzman, J. Frazer, M. Dias, D. Franceschi, R. Orenbuch, Y. Gal, D. S. Marks, ProteinGym: Large-scale benchmarks for protein design and fitness prediction, bioRxivorg (2023). 10.1101/2023.12.07.570727.

38. J. Meier, R. Rao, R. Verkuil, J. Liu, T. Sercu, A. Rives, Language models enable zero-shot prediction of the effects of mutations on protein function, bioRxiv (2021). 10.1101/2021.07.09.450648.

39. P. Notin, M. Dias, J. Frazer, J. Marchena-Hurtado, A. Gomez, D. S. Marks, Y. Gal, Tranception: protein fitness prediction with autoregressive transformers and inference-time retrieval, arXiv [cs.LG] (2022). 10.48550/arXiv.2205.13760.

40. C. Marquet, M. Heinzinger, T. Olenyi, C. Dallago, K. Erckert, M. Bernhofer, D. Nechaev, B. Rost, Embeddings from protein language models predict conservation and variant effects. Hum. Genet. 141, 1629–1647 (2022).

41. E. Nijkamp, J. A. Ruffolo, E. N. Weinstein, N. Naik, A. Madani, ProGen2: Exploring the boundaries of protein language models. Cell Syst. 14, 968–978.e3 (2023).

42. Z. Lin, H. Akin, R. Rao, B. Hie, Z. Zhu, W. Lu, N. Smetanin, R. Verkuil, O. Kabeli, Y. Shmueli, A. Dos Santos Costa, M. Fazel-Zarandi, T. Sercu, S. Candido, A. Rives, Evolutionary-scale prediction of atomic-level protein structure with a language model. Science 379, 1123–1130 (2023).

43. T. Hayes, R. Rao, H. Akin, N. J. Sofroniew, D. Oktay, Z. Lin, R. Verkuil, V. Q. Tran, J. Deaton, M. Wiggert, R. Badkundri, I. Shafkat, J. Gong, A. Derry, R. S. Molina, N. Thomas, Y. A. Khan, C. Mishra, C. Kim, L. J. Bartie, M. Nemeth, P. D. Hsu, T. Sercu, S. Candido, A. Rives, Simulating 500 million years of evolution with a language model. Science 387, 850–858 (2025).

44. A. S. Candido, T. Hayes, A. Derry, R. Rao, Z. Lin, R. Verkuil, B. Z. Wu, J. S. Lee, E. S. Bruguera, J. A. Keval, M. Kopylov, J. E. Pak, W. Wu, N. Thomas, S. Mataraso, A. Hsu, C. Trotman-Grant, K. Fatras, A. dos Santos Costa, R. Badkundri, H. Akin, D. Oktay, J. Deaton, E. Montabana, H. Sitwala, Y. Yu, M. Wiggert, D. A. Carlin, A. W. Goering, T. Blazejewski, M. Sandora, M. Hla, T. Z. Jia, L. H. Kloker, N. J. Sofroniew, M. Uehara, J. Pannu, S. Bachas, D. S. Liu, T. Sercu, A. Rives, Language modeling materializes a world model of protein biology, bioRxiv (2026). 10.64898/2026.06.03.729735.

44. E. Laine, Y. Karami, A. Carbone, GEMME: A simple and fast Global Epistatic Model predicting Mutational Effects. Mol. Biol. Evol. 36, 2604–2619 (2019).

45. T. Bepler, T. Truong Jr, “PoET: A generative model of protein families as sequences-of-sequences” in *Advances in Neural Information Processing Systems 36* (Neural Information Processing Systems Foundation, Inc. (NeurIPS), San Diego, California, USA, 2023), pp. 77379–77415.

46. T. A. Hopf, A. Gazizov, S. Garcia Busto, E. Eschbach, S. Lee, M. Mirdita, R. Orenbuch, K. Belahsen, D. Ross, C. Sander, M. Steinegger, S. d’Oelsnitz, D. S. Marks, Evedesign: Accessible biosequence design with a unified framework, bioRxiv (2026). 10.64898/2026.03.17.712115.

47. K. K. Yang, N. Zanichelli, H. Yeh, Masked inverse folding with sequence transfer for protein representation learning. Protein Eng. Des. Sel. 36, gzad015 (2023).

48. J. Su, C. Han, Y. Zhou, J. Shan, X. Zhou, F. Yuan, SaProt: Protein language modeling with structure-aware vocabulary, bioRxiv (2023). 10.1101/2023.10.01.560349.

49. M. Li, P. Tan, X. Ma, B. Zhong, H. Yu, Z. Zhou, W. Ouyang, B. Zhou, L. Hong, Y. Tan, ProSST: Protein language modeling with quantized structure and disentangled attention, bioRxiv (2024). 10.1101/2024.04.15.589672.

50. M. Tekpinar, L. David, T. Henry, A. Carbone, PRESCOTT: a population aware, epistatic, and structural model accurately predicts missense effects. Genome Biol. 26, 113 (2025).

51. M. Tsishyn, P. Hermans, M. Rooman, F. Pucci, Residue conservation and solvent accessibility are (almost) all you need for predicting mutational effects in proteins. Bioinformatics 41, btaf322 (2025).

52. M. Abakarova, M. I. Freiberger, A. Liehrmann, M. Rera, E. Laine, Proteome-wide prediction of the functional impact of missense variants with ProteoCast. Nat. Commun. 17 (2026).

53. Y. Zhou, Q. Pan, D. E. V. Pires, C. H. M. Rodrigues, D. B. Ascher, DDMut: predicting effects of mutations on protein stability using deep learning. Nucleic Acids Res. 51, W122–W128 (2023).

54. H. Dieckhaus, M. Brocidiacono, N. Z. Randolph, B. Kuhlman, Transfer learning to leverage larger datasets for improved prediction of protein stability changes. Proc. Natl. Acad. Sci. U. S. A. 121, e2314853121 (2024).

55. O. Dutton, S. Bottaro, M. Invernizzi, I. Redl, A. Chung, F. Hoffmann, L. Henderson, L. Henderson, S. Ruschetta, F. Airoldi, B. Owens, P. Foerch, C. Fisicaro, K. Tamiola, Improving Inverse Folding models at Protein Stability Prediction without additional Training or Data, bioRxiv (2024). 10.1101/2024.06.15.599145.

56. C. Hsu, R. Verkuil, J. Liu, Z. Lin, B. Hie, T. Sercu, A. Lerer, A. Rives, Learning inverse folding from millions of predicted structures, bioRxiv (2022). 10.1101/2022.04.10.487779.

57. Y. Zhou, Y. Myung, C. H. M. Rodrigues, D. B. Ascher, DDMut-PPI: predicting effects of mutations on protein-protein interactions using graph-based deep learning. Nucleic Acids Res. 52, W207–W214 (2024).

58. S. Gurev, N. Youssef, N. Jain, A. Mehrotra, S. R. M. Leung, A. Jackson, D. Marks, Evaluating variant effect prediction across viruses, bioRxiv (2025). 10.1101/2025.08.04.668549.

59. M. C. Ekechukwu, D. J. Oberste, B. A. Fane, Host and phi X 174 mutations affecting the morphogenesis or stabilization of the 50S complex, a single-stranded DNA synthesizing intermediate. Genetics 140, 1167–1174 (1995).

60. R. A. Bernal, S. Hafenstein, R. Esmeralda, B. A. Fane, M. G. Rossmann, The phiX174 protein J mediates DNA packaging and viral attachment to host cells. J. Mol. Biol. 337, 1109–1122 (2004).

61. L. N. Young, A. M. Hockenberry, B. A. Fane, Mutations in the N terminus of the øX174 DNA pilot protein H confer defects in both assembly and host cell attachment. J. Virol. 88, 1787–1794 (2014).

62. A. P. Roznowski, J. M. Fisher, B. A. Fane, Mutagenic analysis of a DNA translocating tube’s interior surface. Viruses 12, E670 (2020).

63. M. Chandler, F. de la Cruz, F. Dyda, A. B. Hickman, G. Moncalian, B. Ton-Hoang, Breaking and joining single-stranded DNA: the HUH endonuclease superfamily. Nat. Rev. Microbiol. 11, 525–538 (2013).

64. S. M. Doore, C. D. Baird, 2012 University of Arizona Virology Undergraduate Lab, A. P. Roznowski, B. A. Fane, The evolution of genes within genes and the control of DNA replication in microviruses. Mol. Biol. Evol. 31, 1421–1431 (2014).

65. J. Abramson, J. Adler, J. Dunger, R. Evans, T. Green, A. Pritzel, O. Ronneberger, L. Willmore, A. J. Ballard, J. Bambrick, S. W. Bodenstein, D. A. Evans, C.-C. Hung, M. O’Neill, D. Reiman, K. Tunyasuvunakool, Z. Wu, A. Žemgulytė, E. Arvaniti, C. Beattie, O. Bertolli, A. Bridgland, A. Cherepanov, M. Congreve, A. I. Cowen-Rivers, A. Cowie, M. Figurnov, F. B. Fuchs, H. Gladman, R. Jain, Y. A. Khan, C. M. R. Low, K. Perlin, A. Potapenko, P. Savy, S. Singh, A. Stecula, A. Thillaisundaram, C. Tong, S. Yakneen, E. D. Zhong, M. Zielinski, A. Žídek, V. Bapst, P. Kohli, M. Jaderberg, D. Hassabis, J. M. Jumper, Accurate structure prediction of biomolecular interactions with AlphaFold 3. Nature 630, 493–500 (2024).

66. S. Eisenberg, J. Griffith, A. Kornberg, phiX174 cistron A protein is a multifunctional enzyme in DNA replication. Proc. Natl. Acad. Sci. U. S. A. 74, 3198–3202 (1977).

67. D. R. Brown, T. Schmidt-Glenewinkel, D. Reinberg, J. Hurwitz, DNA sequences which support activities of the bacteriophage phi X174 gene A protein. J. Biol. Chem. 258, 8402–8412 (1983).

68. A. D. van Mansfeld, S. A. Langeveld, P. D. Baas, H. S. Jansz, G. A. van der Marel, G. H. Veeneman, J. H. van Boom, Recognition sequence of bacteriophage phi X174 gene A protein--an initiator of DNA replication. Nature 288, 561–566 (1980).

69. A. D. van Mansfeld, H. A. van Teeffelen, P. D. Baas, H. S. Jansz, Two juxtaposed tyrosyl-OH groups participate in phi X174 gene A protein catalysed cleavage and ligation of DNA. Nucleic Acids Res. 14, 4229–4238 (1986).

70. J. E. Ikeda, A. Yudelevich, N. Shimamoto, J. Hurwitz, Role of polymeric forms of the bacteriophage phi X174 coded gene A protein in phi XRFI DNA cleavage. J. Biol. Chem. 254, 9416–9428 (1979).

71. N. M. Luscombe, R. A. Laskowski, J. M. Thornton, Amino acid-base interactions: a three-dimensional analysis of protein-DNA interactions at an atomic level. Nucleic Acids Res. 29, 2860–2874 (2001).

72. M. H. Werner, A. M. Gronenborn, G. M. Clore, Intercalation, DNA kinking, and the control of transcription. Science 271, 778–784 (1996).

73. J. L. Kim, D. B. Nikolov, S. K. Burley, Co-crystal structure of TBP recognizing the minor groove of a TATA element. Nature 365, 520–527 (1993).

74. S. Korolev, J. Hsieh, G. H. Gauss, T. M. Lohman, G. Waksman, Major domain swiveling revealed by the crystal structures of complexes of E. coli Rep helicase bound to single-stranded DNA and ADP. Cell 90, 635–647 (1997).

75. G. S. Goetz, S. Englard, T. Schmidt-Glenewinkel, A. Aoyama, M. Hayashi, J. Hurwitz, Effect of phi X C protein on leading strand DNA synthesis in the phi X174 replication pathway. J. Biol. Chem. 263, 16452–16460 (1988).

76. D. F. Burke, P. Bryant, I. Barrio-Hernandez, D. Memon, G. Pozzati, A. Shenoy, W. Zhu, A. S. Dunham, P. Albanese, A. Keller, R. A. Scheltema, J. E. Bruce, A. Leitner, P. Kundrotas, P. Beltrao, A. Elofsson, Towards a structurally resolved human protein interaction network. Nat. Struct. Mol. Biol. 30, 216–225 (2023).

77. I. R. Humphreys, J. Pei, M. Baek, A. Krishnakumar, I. Anishchenko, S. Ovchinnikov, J. Zhang, T. J. Ness, S. Banjade, S. R. Bagde, V. G. Stancheva, X.-H. Li, K. Liu, Z. Zheng, D. J. Barrero, U. Roy, J. Kuper, I. S. Fernández, B. Szakal, D. Branzei, J. Rizo, C. Kisker, E. C. Greene, S. Biggins, S. Keeney, E. A. Miller, J. C. Fromme, T. L. Hendrickson, Q. Cong, D. Baker, Computed structures of core eukaryotic protein complexes. Science 374, eabm4805 (2021).

78. O. Follonier, Y. Liu, P. Campomanes, L. Lafrenaye, J. Racle, D. Alvarez, J. van Gerwen, R. Heinzmann, J. Jänes, E. Kummelstedt, J. Durairaj, D. Gfeller, S. Vanni, P. Beltrao, Capabilities, specificity gaps and training-data dependence of AlphaFold3 across diverse application areas, bioRxiv (2026). 10.64898/2026.07.13.738147.

79. M. A. Coelho, M. E. Strauss, A. Watterson, S. Cooper, S. Bhosle, G. Illuzzi, E. Karakoc, C. Dinçer, S. F. Vieira, M. Sharma, M. Moullet, D. Conticelli, J. Koeppel, K. McCarten, C. M. Cattaneo, V. Veninga, G. Picco, L. Parts, J. V. Forment, E. E. Voest, J. C. Marioni, A. Bassett, M. J. Garnett, Base editing screens define the genetic landscape of cancer drug resistance mechanisms. Nat. Genet. 56, 2479–2492 (2024).

80. M. G. Schubert, E. Sukarto, F. R. Martin, J. E. Mancuso, A. Spens, T. Pedersen, K. Isaev, H. Greene, N. D. Hicks, U. Nattermann, J. Liu, D. A. Stork, E. A. DeBenedictis, BioBloom, a method for barcoded saturation mutagenesis of an entire bacterial genome, bioRxiv (2026). 10.64898/2026.02.04.703572.

81. S. Li, S. C. Vonesch, K. R. Roy, C. S. Tu, F. Steudle, M. Nguyen, C. Jann, L. M. Steinmetz, The editable landscape of the yeast genome reveals hotspots of structural variant formation. Sci. Adv. 11, eady9875 (2025).

82. P. M. Sharp, W. H. Li, The codon Adaptation Index--a measure of directional synonymous codon usage bias, and its potential applications. Nucleic Acids Res. 15, 1281–1295 (1987).

83. A. J. Faure, J. M. Schmiedel, P. Baeza-Centurion, B. Lehner, DiMSum: an error model and pipeline for analyzing deep mutational scanning data and diagnosing common experimental pathologies. Genome Biol. 21, 207 (2020).

84. E. C. Meng, T. D. Goddard, E. F. Pettersen, G. S. Couch, Z. J. Pearson, J. H. Morris, T. E. Ferrin, UCSF ChimeraX: Tools for structure building and analysis. Protein Sci. 32, e4792 (2023).

85. T. A. Hopf, A. G. Green, B. Schubert, S. Mersmann, C. P. I. Schärfe, J. B. Ingraham, A. Toth-Petroczy, K. Brock, A. J. Riesselman, P. Palmedo, C. Kang, R. Sheridan, E. J. Draizen, C. Dallago, C. Sander, D. S. Marks, The EVcouplings Python framework for coevolutionary sequence analysis. Bioinformatics 35, 1582–1584 (2019).

86. M. A. Stiffler, D. R. Hekstra, R. Ranganathan, Evolvability as a function of purifying selection in TEM-1 β-lactamase. Cell 160, 882–892 (2015).

87. D. H. Mathews, M. D. Disney, J. L. Childs, S. J. Schroeder, M. Zuker, D. H. Turner, Incorporating chemical modification constraints into a dynamic programming algorithm for prediction of RNA secondary structure. Proc. Natl. Acad. Sci. U. S. A. 101, 7287–7292 (2004).

